# 3D pose estimation enables virtual head-fixation in freely moving rats

**DOI:** 10.1101/2022.04.14.488303

**Authors:** Artur Schneider, Christian Zimmermann, Mansour Alyahyay, Florian Steenbergen, Thomas Brox, Ilka Diester

**Affiliations:** Optophysiology Lab, Institute of Biology III, Albert-Ludwigs-University, 79110 Freiburg, Germany; Department of Computer Science, Albert-Ludwigs-University, 79110 Freiburg, Germany; IMBIT//BrainLinks-BrainTools, Albert-Ludwigs-University, 79110 Freiburg, Germany; Bernstein Center Freiburg, Albert-Ludwigs-University, 79110 Freiburg, Germany; CIBSS Centre for Integrative Biological Signalling Studies, Albert-Ludwigs-University, 79110 Freiburg, Germany

**Keywords:** marker-free 3D movement tracking, virtual head-fixation, extracellular recordings, RFA, CFA, motor cortex, optogenetics, tuning curves

## Abstract

The impact of spontaneous movements on neuronal activity has created the need to quantify behavior. We present a versatile framework to directly capture the 3D motion of freely definable body points in a marker-free manner with high precision and reliability. Combining the tracking with neural recordings revealed multiplexing of information in the motor cortex neurons of freely moving rats. By integrating multiple behavioral variables into a model of the neural response, we derived a virtual head-fixation for which the influence of specific body movements was removed. This strategy enabled us to analyze the behavior of interest (e.g., front paw movements). Thus, we unveiled an unexpectedly large fraction of neurons in the motor cortex with tuning to the paw movements, which was previously masked by body posture tuning. Once established, our framework can be efficiently applied to large datasets while minimizing the experimental workload caused by animal training and manual labeling.

## 1. Introduction

Systems neuroscience strives to assign functions to neuronal circuits and their activity. Often, these functions are defined by behavior in a task designed to cover specific aspects, such as rule learning or reaching to a defined target. However, neuronal activity can be dominated by uninstructed movements not required for the task. Confusingly, some uninstructed movements can be aligned to trial events, whereas other movements might occur idiosyncratically, accounting for trial-by-trial fluctuations that are often considered “noise” (Musall et al., 2019). It has become increasingly clear that a considerable part of this neural variability is due to movement signals that were not previously measured and accounted for (Parker et al., 2020). These uninstructed movements strongly influence task-related activity even in studies conducted with head-fixed animals engaged in well-controlled behavioral paradigms (Stringer et al., 2019; Salkoff et al., 2019; Musall et al., 2019; Allen et al., 2017). Additionally, information processing in the brain, particularly in cortical areas, occurs in a parallel manner (Ledberg et al., 2007; Stringer et al., 2019; Steinmetz et al., 2019). Thus, our interpretations of brain function in highly isolated environments might be impacted. To alleviate this problem, a more voluntary, freely moving approach to behavior has been suggested (Cisek and Kalaska, 2010; Parker et al., 2020). Highly reductionist experimental paradigms, which rely on head-fixation and instructed, taught movements, can also account for movements that occur out of the task context (Stringer et al., 2019; Salkoff et al., 2019; Musall et al., 2019). However, in head-fixed experiments, the disruption of the naturalistic movement patterns influences the interpretation of the movement-related signals (Parker et al., 2020).

To address these challenges, we inverted the classical research approach and asked whether it is possible to relate neuronal activity to movements of individual body parts in freely moving animals that conducted only spontaneous movements. For this approach, two requisites must be met: (1) detailed 3D tracking of the animals’ unconstrained movements at the level of single body parts; (2) isolation of the body part of interest from the influence of other body parts. Such an approach would enable analyses of instructed as well as uninstructed movements with unprecedented precision and from an entirely new scientific perspective (i.e., with a focus on individual body parts integrated into full body movements). It would also reduce the training time of animals due to the lack of constraints, and bring neuroscientific findings from an artificial laboratory setting closer to a more naturalistic scenario.

New video tracking tools have augmented the scope of experiments in recent years. However, existing tools are still limited by at least one of the following issues. First, marker-based approaches (Mimica et al., 2018; Marshall et al., 2021) can influence natural movements, are restricted to applicable body sites, and rely on the animal’s tolerance. Second, thus far, marker-free analyses have mostly been applied in 2D settings (Graving et al., 2019; Mathis et al., 2018; Pereira et al., 2019), which do not pose the need for detailed pose reconstruction of freely moving animals covering all three spatial dimensions. When recording with multiple cameras, 2D outputs can be triangulated to reconstruct a 3D pose a posteriori (Günel et al., 2019; Mathis and Mathis, 2019; Karashchuk et al., 2021; Nath et al., 2019), but such post-processing suffers from the ambiguities in the initial 2D analysis, reducing both accuracy and reliability (Fig. 1**b** left).

**Figure 1:**
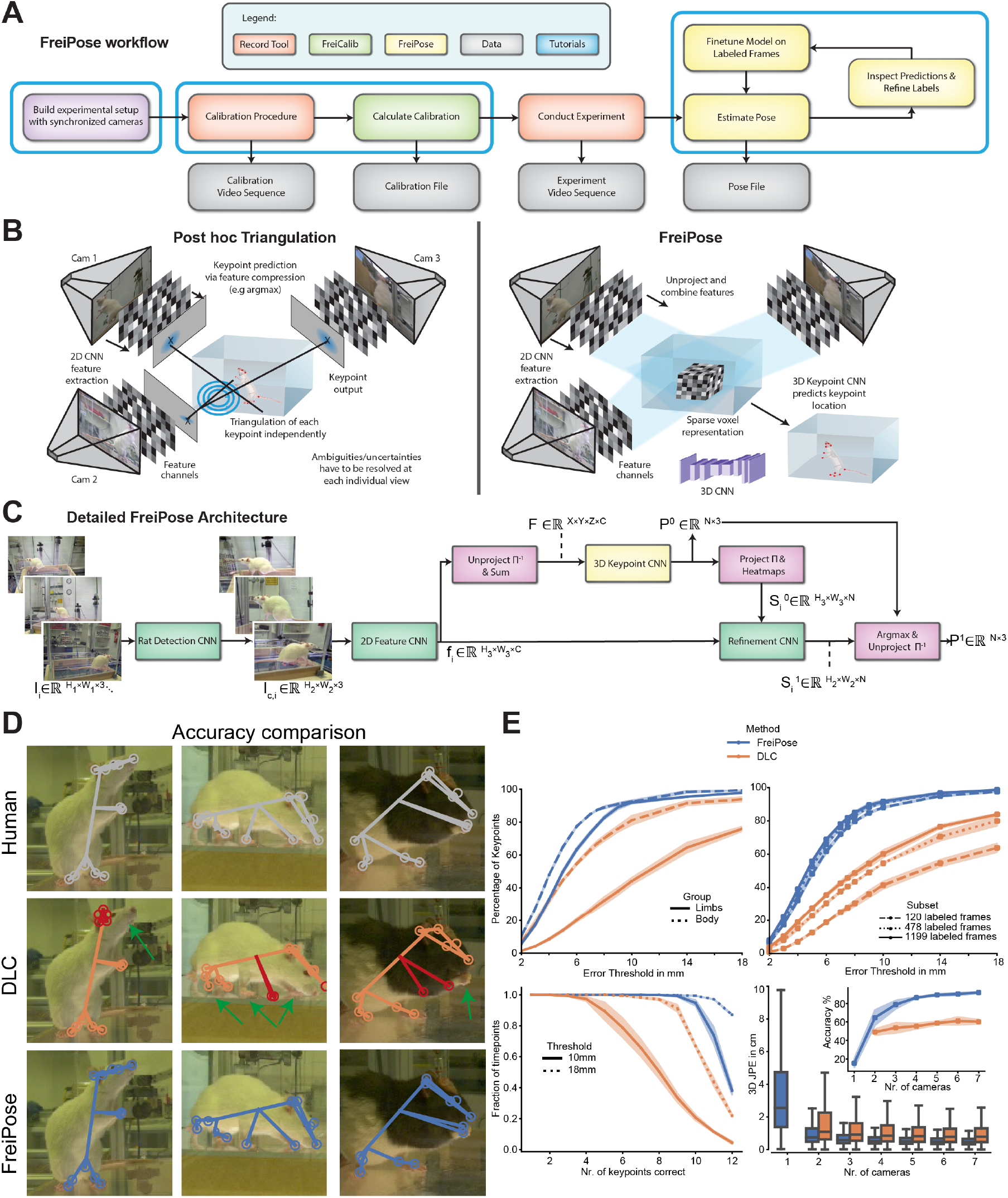
Overview of the motion capture framework and conceptual comparison with two existing approaches. **a** Workflow of an experiment based on FreiPose. Temporally synchronized cameras are calibrated using FreiCalib and RecordTool. The pose is estimated from recordings in a predict-inspect-refine loop, where predictions can be refined if they are not yet sufficiently accurate. **b** Commonly used motion capture approaches (e.g., DLC) reconstruct 2D poses for each view independently and subsequently calculate a 3D pose (left). This post hoc triangulation requires the resolution of ambiguities in each view separately, and errors are carried over to the 3D triangulation. FreiPose accumulates evidence from all views (via projection of features into common 3D space), leading to a holistic 3D pose reconstruction (right). Ambiguities are resolved after information from all views becomes available. **c** FreiPose architecture. A bounding box network is applied to the camera images. Afterwards, image features are extracted from the cropped images and unprojected into a common 3D representation. This 3D representation is used to reconstruct the initial pose ***P*** ^0^, which is projected into the views for further refinement. Finally, the refined 2D reconstructions are used to calculate the final 3D pose ***P*** ^1^ (Zimmermann and Brox, 2017). **d** Qualitative accuracy comparison. Three representative examples with projected predicted keypoints are shown. FreiPose predictions (blue) were highly similar to the human labels (grey), whereas the DLC predictions (orange) for some keypoints were visibly wrong (indicated by red markers as well as green arrows). **e** Quantitative accuracy comparison. Top-left: Percentage of correct keypoints within threshold (PCK) for FreiPose (blue) and DLC (orange) for body as well as limb keypoints. Note the substantial reduction in accuracy for DLC for the highly articulated limbs. Top-right: PCK for networks using a subset of the entire dataset to indicate data efficiency. Bottom-left: Fraction of time with keypoints within threshold. Bottom-right: 3D joint position error (JPE) and accuracy (insert, 10 mm threshold) depending on the number of cameras used for training. FreiPose reached high accuracy with only 3–4 cameras and consistently outperformed DLC.

To address these challenges, a novel approach (DANNCE) (Dunn et al., 2021) has been suggested. DANNCE allows a 3D convolutional neural network (CNN) to infer landmark locations using information across cameras. Using a large marker-based dataset for rats, Dunn et al. were able to demonstrate the advantages of native 3D keypoint prediction compared to 2D methods such as DeepLabCut (DLC) (Mathis et al., 2018). DANNCE uses a volumetric approach whereby the dense image representation is unprojected into common 3D space (Fig. S. 1**a** middle).

We developed a marker-free, animal tracking tool with native 3D predictions, which we refer to as FreiPose. This tool allows us to reconstruct detailed body poses and single body part movements in freely moving animals directly in 3D. From synchronized multi-view videos, FreiPose reconstructs a flexible set of on-body keypoints, which can be defined freely in terms of number and position on the animal (eye, paw, etc.). The novelty of FreiPose is the projection of learned, sparse image features from individual camera views into a common 3D space, where a 3D CNN is able to combine all information to reason about keypoint location (Fig. 1**b** right). This approach is different from DANNCE, which projects volumetrically processed full images rather than sparse learned features. Due to its multi-view approach and native 3D reconstruction, FreiPose is particularly suitable for freely moving animals in various environments, including obstructions. It is a versatile tool, which can be applied flexibly in multiple experimental settings. For instance, we used it for full-body tracking of rats, mice, cowbirds, and marmoset, as well as for detailed tracking of rat paws, including individual digits. Furthermore, we provide a full pipeline for the workflow, including recording, triangulation, and labeling tools.

A second prerequisite for extracting the neuronal representations of movements of individual body parts relies on the reduction of the contributions of other body parts and movements. In freely moving rats, we found a small subset of neurons to be tuned to single paw trajectories similar to what has been described previously in head-fixed animals engaged in constrained tasks (Kakei et al., 1999; Moran and Schwartz, 1999). Considering the parallel, multiplexed information processing in the cortex, we introduced a “virtual head-fixation” approach to account for the movement and behavioral variations that occur during freely moving experiments. This approach relies on statistical modeling of the simultaneous influence of individual movement variables captured by FreiPose onto single unit activity. We employed non-parametric regression in the form of generalized additive models (GAM) (Hastie and Tibshirani, 1990), which combine the flexibility of non-parametric modeling with the interpretability of linear models. Based on a fitted model for an individual neuron, the influence of defined movement variables (uninstructed movements e.g., head movements) can be removed. This approach revealed that far more neurons in the motor cortex were tuned to paw movements than has been previously described (Isomura et al., 2009), indicating that the movements of other body parts masked the paw movement encoding.

## 2. Results

We developed a full pipeline (Fig. 1**a**) for the native 3D prediction of anatomical landmarks (keypoints) on animals based on a multi-camera approach. Model training follows an iterative three-step reconstructinspect-refine scheme (Fig. 1**a**): the pre-trained network computes pose reconstructions of the animal in the video. Guided by reconstruction confidence, potentially erroneous frames are selected for manual correction in a human-in-the-loop approach (see Fig. S. 9**a**). Multiple persons can annotate videos in parallel using a standalone, intuitive drag-and-drop GUI that leverages multi-view constraints during annotation (see Fig. S. 9**b**). The method is retrained with the additional annotation, leading to pose reconstructions with fewer and smaller errors. Once the network is sufficiently trained, it produces 3D poses without further manual intervention.

FreiPose comes with a *Docker* installation package, which simplifies the software installation and tutorials with exemplary data and step-by-step videos. All necessary information for applying FreiPose can be found in the released GitHub repositories.

FreiPose leverages an experimental setup with multiple cameras (Fig. 1**b**) and a dedicated convolutional network architecture to detect and track predefined 3D keypoints on the animal’s body. This approach integrates information across all available views (Fig. 1**b** right) instead of processing views separately (Fig. 1**b** left). Indeed, due to occlusion, amongst other things, it is often impossible from a single view to measure the precise location of all keypoints, yet existing 2D methods (Fig. 1**b** left) attempt to reconstruct the keypoint locations in the images. This is problematic in case of ambiguities, for example, between the left and the right paw. If the left paw is not visible in some views, DLC will detect the keypoint on the right paw with high confidence. It is hard to impossible to fix these mistakes in the post hoc triangulation. Such shortcomings lead to diminishing performance in the subsequent post hoc triangulation step (see Fig. 1**d** and Fig. S. 1**b**). In freely moving animals, occlusions are almost unavoidable due to the rich repertoire of movements, which underlines the importance of a 3D approach. FreiPose extracts features rather than keypoints from the images and deploys an inverse projection (Zimmermann et al., 2019), which maps features into a 3D representation and deploys a 3D CNN (Falk et al., 2019) on the voxelized representation (Fig. 1 **b** right and **c**). FreiPose learns to reason on the joint representation and reconstructs a 3D pose incorporating information from all views.

### FreiPose captured motion with high accuracy and reliability

To test the feasibility of FreiPose to track rodent movements, we let rats roam freely in a box without any specific task. We measured the accuracy (median 3D error) and reliability (percentage of samples below a maximum error bound) of FreiPose on video recordings of the rats consisting of 1813 manually labeled samples with 12 distinct body keypoints (see Fig. S. 8). The frames were sampled from 12 recording sessions featuring 5 different individuals (3 Long-Evans (hooded) and 2 Sprague Dawley (albino) rats). We split the recordings into training and evaluation sets, with 1199 training samples and 614 evaluation samples. Each sample contained 7 images recorded simultaneously from different cameras and a single, manually annotated 3D pose consisting of a 3D location for each keypoint which was obtained from manual 2D annotation in at least two camera views.

For comparison, we trained DLC, a popular tool for 2D keypoint tracking, on the same dataset of images and applied robust triangulation methods to yield 3D poses. During visual qualitative comparison, it became clear that DLC struggles with partially occluded keypoints, which are then incorrectly triangulated (see Fig. 1**d** and Fig. S. 1**b**). Quantitatively, FreiPose compared favorably in terms of accuracy (median error of 4.54 mm vs. 7.81 mm for the full sample setting), particularly for the highly articulated limbs (Fig. 1**e** top and Fig. S. 1**c**). FreiPose also outperformed DLC in terms of reliability (percentage of keypoints with an error smaller than 10 mm was 91.8% vs. 60.2%; furthermore, 75% of time points *>*= 10 keypoints were within 10 mm threshold, whereas only 11.8% for DLC Fig. 1**e**, lower left). Additionally, FreiPose was more data efficient (lower median error with the same number of labeled samples, Fig. 1**e** upper right), and required fewer camera views to reach a certain level of accuracy (Fig. 1**e** lower right).

### FreiPose can be adapted to different tasks and species

The need for accurate movement quantification extends beyond rodents. As such, FreiPose was created without any species bias and can generalize to any animal or object. Thus, it was easy to adapt FreiPose to new experimental setups with a highly distinct skeleton shape, such as tracking of a rat’s paw including digit segments during pellet reaching (0.32 mm median error Fig. 2**a,b**). We then extended our experiments to different species such as cowbirds in a large (2.5 m × 2.5 m × 6 m) enclosure (Badger et al., 2020) (2.92 cm median error Fig. 2**c**,**d**), a marmoset in an enriched homecage (Dunn et al., 2021) (2.6 cm median error Fig. 2**e**,**f**), and a miniscope-wearing mouse (3.5 mm median error Fig. 2**g**,**h**). For the adaptation to the new settings, only small sets of labelled frames (e.g., 78 for detailed paw tracking Fig. 2**a**,**b**) were necessary to obtain a very good performance. For more details on the datasets, see Methods and Fig. S. 2.

**Figure 2:**
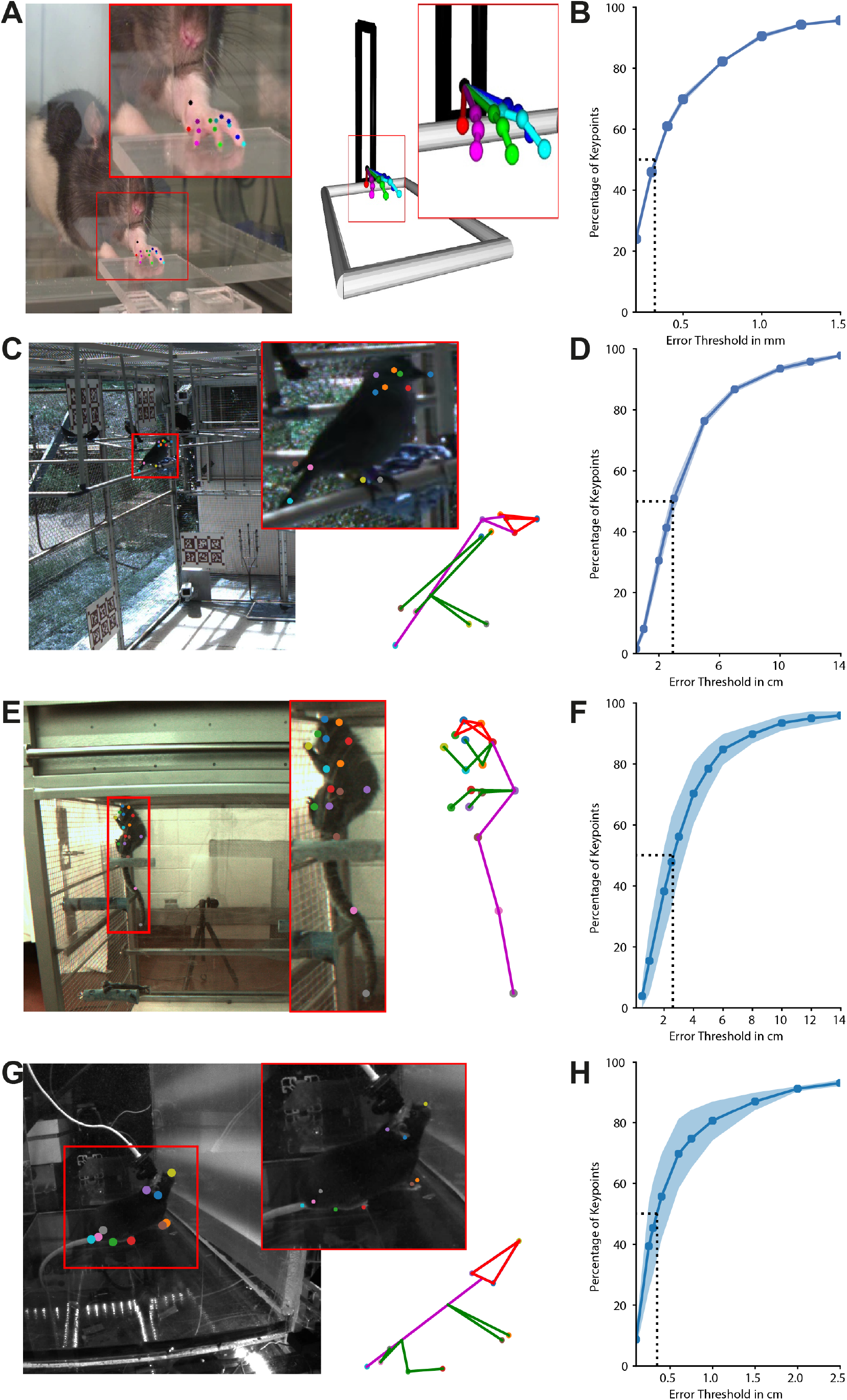
FreiPose can be easily adapted towards other species and tasks. Exemplary predictions on different datasets. Shown is a single camera view, zoomed in view (left), as well as a skeletal representation (right) of predicted 3D keypoints. **a** Paw reconstruction including individual digits during a pellet-reaching task. **c** Cowbird dataset, **e** Marmoset dataset, and **g** Mouse dataset. **b, d, f, h** PCK plots for corresponding species. Shaded areas indicate SEM for 3-fold cross-validation. Dotted lines indicate median error (referring to 50% of the PCK).

### FreiPose tracking enabled behavioral categorization

The rich information regarding the animal’s motion and posture obtained by FreiPose provides a range of opportunities for higher-level holistic as well as profound kinematic analysis of individual movements. Combining the temporal aspects of multiple keypoint positions, we aimed to identify the concurrent behavior of the animal. To this end, we developed a supervised machine learning approach for the segmentation of behavioral categories. We then applied this algorithm to extract behavioral motifs, as well as to exclude defined behavioral categories from the analysis (e.g., grooming) because these behaviors often lead to artifacts in electrophysiological measurements. Based on keypoint representations from FreiPose, we were able to split test-set videos of freely roaming rats into segments according to behavioral categories in an automated fashion (Fig. 3**b**). For the behavioral classifier, we calculated postural decompositions based on the output of FreiPose. The resulting multidimensional data were embedded into low-dimensional spaces using UMAP, a non-linear dimensionality reduction technique attempting to preserve the global data structure (McInnes et al., 2020). A supervised support vector machine (SVM) classifier was trained on this embedding, using sparsely labeled annotations. We defined 5 distinguishable behavioral clusters: locomotion, grooming, rearing, sitting, and exploration of the nearby environment defined by small sidewards steps and sniffing movements. The SVM classifier recalled the correct classes on the test videos with high fidelity (83% balanced accuracy on test sessions, see Fig. 3**c**). As an alternative approach for cases in which behavioral categories are not known preemptively, we established unsupervised behavioral categorization with FreiPose, leading to the detection of intuitive behavioral clusters (see methods section and Fig. S. 3**b**). We then combined FreiPose–enabled behavioral categorization with simultaneous neural recordings in rats motor cortex (see Methods for details). In accordance with previous studies (Omlor et al., 2019), we demonstrated that the majority (77%, 126*/*164) of neurons had a significantly different firing rate during distinct behaviors, as identified via our supervised approach (see examples in Fig. S. 3**a**), demonstrating the validity of the tracking and the resulting behavioral categorization.

**Figure 3:**
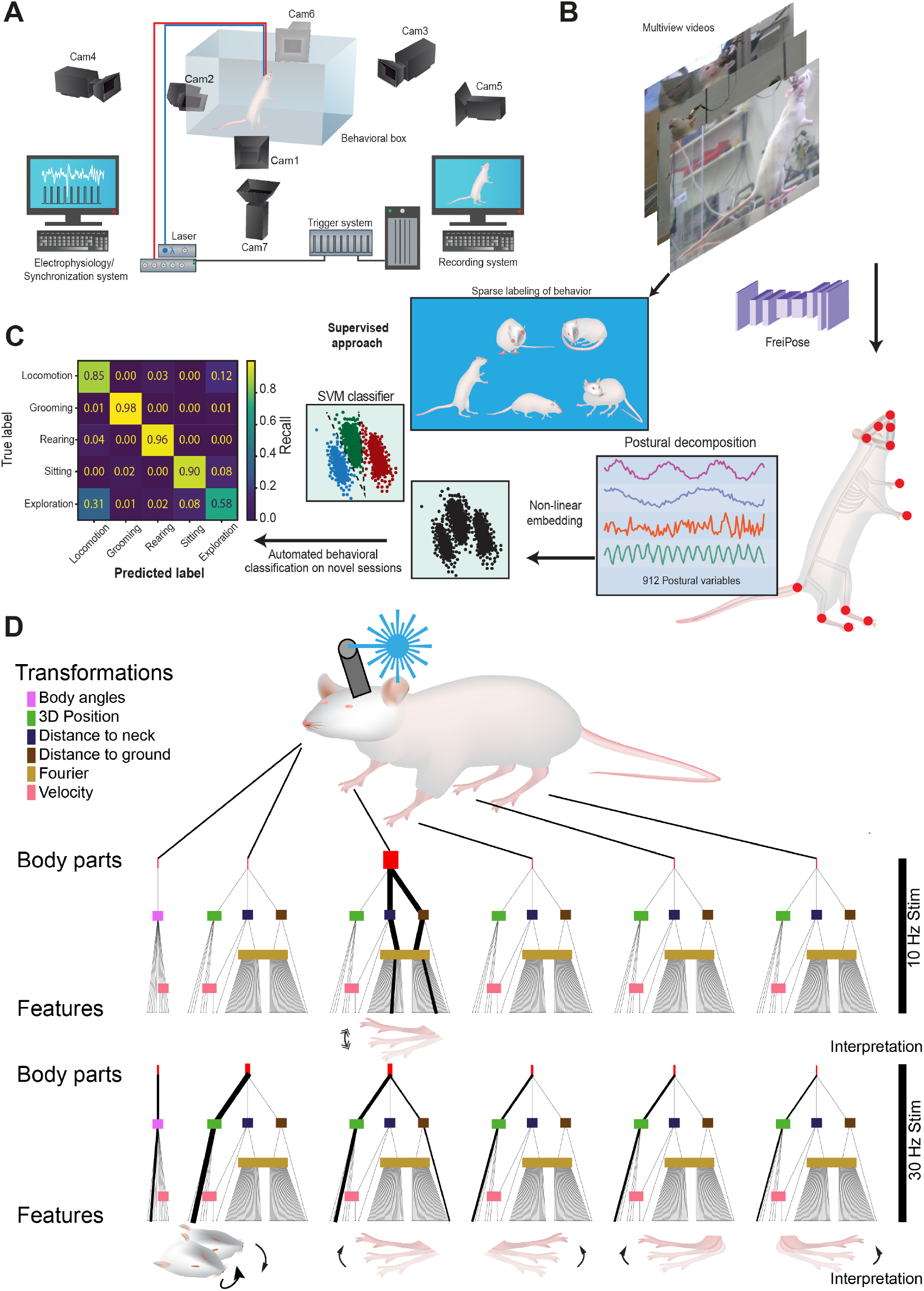
FreiPose allows automated behavioral classification, detection of optogenetic effects on movements. **a** Motion capture setup, compatible with electrophysiology, optogenetic as well as calcium imaging in freely moving animals. **b** Schematic of the automated behavioral classification. Predicted keypoints were embedded into low dimensional space, and a classifier was trained on predefined behaviors. **c** Confusion matrix with results of the supervised behavioral classifier on test session. **d** Automatic detection of optogenetic impact on movements. Attribution of stimulation effect to individual body parts. For each keypoint, a set (n=912) of features was calculated using different transformations (see methods). Ensemble of classifiers identified a subset of important features, which can be back-attributed to the corresponding keypoints. Each line indicates the lineage path of a feature. For significant features, the line width was doubled. Thicker lineage lines indicate multiple significant downstream features. Shown are the combined results of the cross-animal validation.

### FreiPose detected optogenetic stimulation effects on individual body parts

We hypothesized that FreiPose would be capable of detecting deviant behaviors caused by e.g., optogenetic stimulation. This capability would be particularly useful in control studies of optogenetic experiments because it is often required to test for unintended influences of the optogenetic stimulation (Aravanis et al., 2007; Gradinaru et al., 2009; Sun et al., 2021). To test whether FreiPose is suitable for detecting overt behavior during optogenetic stimulation, we injected rats unilaterally with a ChR2-carrying viral vector in the front paw area of the motor cortex. After an expression time of 4 weeks, the rats were stimulated with pulsed (10 or 30 Hz with 10 % duty cycle) blue laser light for 5 s during free exploration of the behavioral box.

Using the behavioral decomposition provided by FreiPose, we identified and attributed the effect of optogenetic stimulation in the motor cortex to the perturbed kinematics. A SVM classifier was able to detect the effect of the optogenetic stimulation (balanced average accuracy of 59.1%-73.1%) and identified the temporal dynamics with an increasingly visible effect over time (Fig. S. 4**a**). To attribute the stimulation effect to individual body parts, we analyzed the classifiers, trained with only a single behavior variable as the input to distinguish important factors from less important factors. The identified features were semantically back-traced to the corresponding origin keypoint (Fig. S. 4 **b**). Using this approach, we were able to identify a rhythmic motion of the contralateral paw during 10 Hz optogenetic stimulation. During the 30 Hz stimulation, the identified motion was relatively tonic and involved the whole body Fig. 3**d**. This is in line with findings in the macaque motor cortex, where increased stimulation frequency led to stronger EMG activation (Watanabe et al., 2020). Additionally, our method generalized across animals, revealing the robust detection capabilities of the approach.

### Body pose reconstruction recapitulated neuronal tuning

Encouraged by the ability to not only identify the general overt behavior during optogenetic stimulation with FreiPose but also to describe the perturbed kinematics, we continued with more detailed analyses of individual paw movements as well as postural changes. Previous work on neuronal representations of body poses in rodents relied on marker-based systems (Hughes et al., 2019; Mimica et al., 2018). Combining FreiPose with simultaneous neural recordings revealed that approximately half of the recorded neurons in the motor cortex were tuned to postural features (50.6%, 83/164 cells Fig. 4**a** and Fig. S. 5**a**,**b**). Neurons were considered tuned if the z-score exceeded a critical value (corresponding to a Bonferroni-corrected p-value < 0.05) relative to the shuffled distribution for at least 3 consecutive postural bins. This fraction of tuned neurons is in accordance with the previously reported percentage of cells in the motor cortex (Mimica et al., 2018), which demonstrates that FreiPose can effectively capture the poses with a precision that allows one to make claims about neuronal correlations.

**Figure 4:**
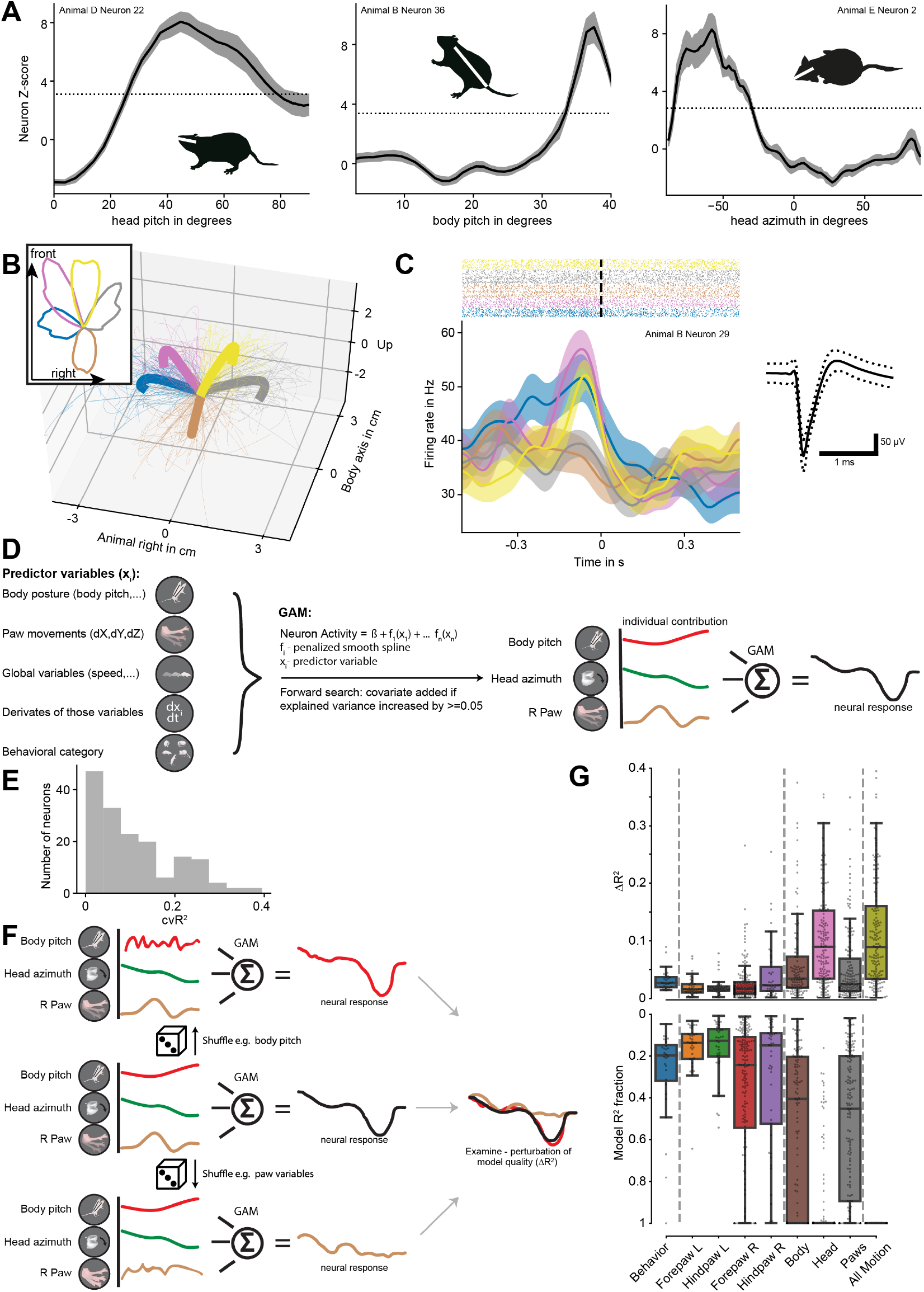
GAM reveals information multiplexing. **a** 1D tuning curves of representative neurons for body pose variables (body pitch, head pitch, and head azimuth) shown as z-score relative to the randomly time-shifted data. Rat pictographs refer to the preferred pose of the respective neuron. Dotted lines indicate the significance threshold, Bonferroni-adjusted z-score, and shaded areas indicate SEM. **b** Trajectories of the right (contralateral) paw. Mean trajectories for each class, as well as a small subset of trajectories, are shown for visualization purposes. Color represents the investigated forepaw directions in the rat’s body reference frame. Inserts indicate the distributions of the horizontal projections of movement directions. Colors represent individual clusters defined by k-means. **c** Representative example neuron with a modulation of the neuronal response relative to the movements of the right paw. Time point 0 represents the movement onset defined by a velocity threshold. Colors indicate the trajectory direction class as defined in panel b and shaded areas indicate SEM. Insert shows the waveform of the corresponding unit ±STD. **d** Schematic of the forward search of the variables to be included in the model. Variables from the list of prospective predictors are added to the model if the new model has higher explained variance. **e** Distribution of explained variance of GAM of all analyzed neurons. **f** Illustration of the approach to estimate the degree of contribution (Δ*R*^2^) of individual variables or groups of variables to the entire model. **g** Explained variance of individual variable groups. Shown is Δ*R*^2^ and its fraction of the full model’s explained variance. The boxes indicate the first and third quartiles. The median is represented by a horizontal line. Whiskers represent 1.5 x IQR. Individual data points (neurons) are represented as dots.

### FreiPose identified neuronal representations of paw trajectories

To test whether we can identify a systematic neuronal representation of spontaneous movements outside of the classical instructed tasks, we extracted single paw trajectories. We divided continuous paw movements into artificial trials by setting a threshold for the paw velocity (see Methods). To validate the manually chosen threshold, we systematically varied it and compared the corresponding population responses. In the range of log_2_(−0.66 to 0.33) no difference in the detected population response was observed (p-value *>* 0.3, one-way ANOVA, see Fig. S. 5 **e**). We clustered paw trajectories by k-means into classes that primarily represented different movement directions (Fig. 4**b**). We found that a small fraction of neurons significantly changed their activity depending on the movement directions (17.68%, 29*/*164, one-way ANOVA of the sum of spikes within ± [333 ms] window around movement onset, with trajectory class as the main effect, per neuron Bonferroni-adjusted p< 0.05, Fig. 4**c** and Fig. S. 5**c**). This result suggests that a subset of neurons in the motor cortex encodes the paw movement trajectory even in a freely moving condition and that the paw trajectory can be reliably tracked with FreiPose.

### Modeling the neuronal response revealed multiplexing

Neurons in the motor cortex contained information not only about the paw movements but also about the behavioral state and body posture, suggesting that cortical neurons multiplex multiple sources of information. Interestingly, neurons often contained information regarding multiple postural variables (see Fig. S. 5**b**). We hypothesized that the variety of behavioral variables that can be captured with FreiPose could help explain the variability of the observed neuronal responses. To analyze the contribution of different behavioral variables, we employed GAM, which provided an unbiased model-based approach to detect the multiplexing of the simultaneous representations of different behavioral variables in individual neurons or neural populations. GAMs reduced the impact of correlated and (partially) equivalent features and, thus, we considered it to be superior to the previously applied bootstrapping approach (based on time-shifted, relative to behavior, recorded neural data). The GAM concept is based on estimated smooth relationships between individual predictors (behavioral variables e.g., body pitch angle) and neural firing rate as the response variable. The smoothness is achieved by penalizing the 2. derivative, which also strongly reduces over-fitting. GAM estimates multiple smooth relationships simultaneously via linear combinations. We established the choice of variables included in the model of an individual neuron via a forward-search procedure (Fig. 4**d**). For a toy example see Fig. S. 10. An example of a predicted firing rate using a GAM is shown in Fig. S. 6**a** and distribution of the explained variance achieved by the models in Fig. 4**e**.

To validate the results of the GAM, we compared the resulting tuning curves for body postures that we obtained during simultaneous fitting of multiple variables with the tuning curves obtained using the bootstrap approach (Fig. 4**a**). 83%, 77% and 79% of neurons with tuning detected via bootstrap for body pitch, head pitch, and head azimuth, respectively, were also found to be modulated using the GAM approach (visual comparison of tuning curve examples in Fig. S. 6**c**). Furthermore, the comparison of the tuning curves detected by the two methods on a population level provided highly similar results (Fig. S. 6**d**).

With the GAM approach, we found that multiple variables were represented at the level of single neurons in the motor cortex. The most commonly represented variables were body posture variables, whereas paw-movement variables were represented less commonly. Additionally, GAM identified not only paw variables of the contralateral paw but also, to a lesser degree, from the ipsilateral side. This finding indicates that the rat motor cortex does not strictly separate signals from contra- and ipsilateral paws, which is similar to findings in monkeys (Ames and Churchland, 2019).

To analyze the contributions of different groups of variables to the model’s explanatory power, we calculated the corresponding Δ*R*^2^ values. The fitted GAM was re-evaluated with a subset of temporally shuffled variables from individual groups. The decrease in the explained variance (Δ*R*^2^) indicates the contribution of those variables (Fig. 4**f**). The variables related to the head and body postures were not only represented in more neurons but also contributed more strongly to the models (0.10 and 0.06 mean Δ*R*^2^ respectively), which is not self-evident because we recorded neuronal signals from a front paw motor area. Paw contribution was 0.05 mean Δ*R*^2^. Total motion was represented in the neural firing rate to a similar degree as that previously described for uninstructed movements during head-fixation (0.11 ± 0.09 Δ*R*^2^ all motion variables (mean±std) vs 0.212 ± 0.012 Δ*R*^2^ in Musall et al. (2019), Fig. 4**g**).

### Virtual head-fixation unmasked a large fraction of paw-tuned neurons

The results from the GAM analysis encouraged us to further investigate the relationship of the multiplexed neuronal representation of paw movements and body postures. Because neurons carried a mixture of information sources, the variability of body postures during freely moving behavior might impact our ability to detect the isolated neural signal of paw movements.

Given that GAM aims to achieve the best possible explanation of the neuronal firing rate based on available movement information obtained via FreiPose, we calculated a hypothetical neural response under “head-fixed” conditions. The influence of body posture variables was removed by fixing their value in the model to their session median. Thus, we were able to create a “virtually head-fixed” condition in which only the impact of the paw movements on neuronal activity remained (Fig. 5**a**).

**Figure 5:**
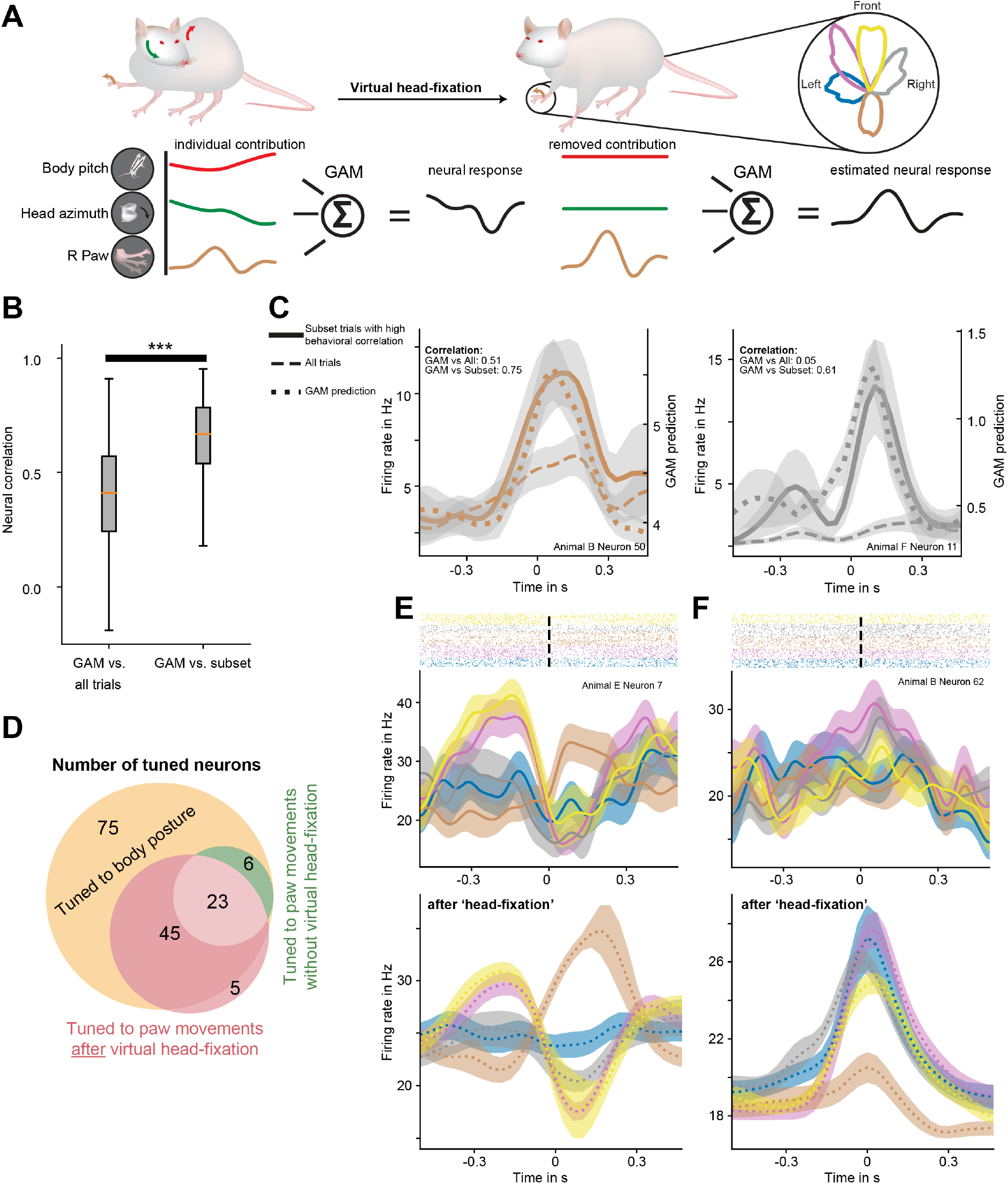
“Virtual head-fixation” describes neural activity during postural rest. **a** Schematic of the virtual head-fixation approach. **b** Here, we contrasted paw movements from all trials in a session, a subset of paw movements with low postural variability with corresponding GAM “head-fixed” predictions. On average, across all neurons and trajectory classes, the GAM predictions had a significantly stronger correlation to the neural response during the subset of trials than to the response during the entire session. *** indicate p< 0.001, Wilcoxon signed-rank test. **c** Representative individual examples for neuronal response to certain trajectory classes. Here, we contrast the average over all trials in a session (dashed line) with the subset of paw movements with low postural variability (filled line). GAM’s “head-fixed” prediction for the subset is shown as a dotted line. The colors represent the trajectory class, and shaded areas represent SEM. The GAM predictions were more similar to the neural response during the subset of trials with low postural variability than the response during all trials, indicating that the GAM “head-fixed” prediction is a valid description of the neural responses during low postural change. **d** Venn diagram displaying number of neurons detected as tuned for paw motion using both “classical” and “head-fixed” approaches as well as posture tuned as identified via GAM. Note that the six neurons not detected using the “head-fixed” approach were likely due to the strong contribution of potentially correlated body posture tuning. **e, f** Representative example neurons. Measured response to the trajectory class versus isolated predicted response after virtual head-fixation. Measured neural response is indicated by filled line and “head-fixed” prediction by dotted line.

To validate this approach, we used a subset of our data, in which the animal executed some paw movements while holding its body mostly still. On average, 21.1 ± 0.6 (%±SEM) of defined paw movements occurred during low postural variability. We argue that a biologically plausible virtual head-fixation would result in more similar predictions during movement periods in which the body posture was stationary than with paw movements that occurred simultaneously with other body movements. In this first scenario, the animal moves its paws but not its body, generating a setting that is similar to movements occurring during virtual head-fixation. We defined the measure for the resemblance of the behavior “behavioral correlation” and the measure for the resemblance of the neural responses “neural correlation” (see Methods). We analyzed whether the GAM-predicted “head-fixed” response was more similar to the firing rate of neurons during a subset of paw movements than to the firing rate over all trials in a session. We define this subset in a manner that trials had a high behavioral correlation to the “head-fixed” behavior. During those trials, only low postural variations occurred. Indeed, the neural response in these trials differed from the average across all trials and was more similar to the GAM prediction (0.41 vs 0.67 median neural correlation, p< 0.001, Wilcoxon signed-rank test) (Fig. 5**b**). This result indicates that the virtual head-fixation approach is a reasonable description of the isolated neural response to the paw movements. We confirmed this finding on a neuronal level by identifying individual neurons with response patterns during low postural variation trials, which strongly resemble head-fixed predictions in contrast to response patterns averaged across all trials (Fig. 5**c**, Fig. S. 7**a**).

By applying the “virtual head-fixation” approach to the entire dataset, we uncovered a large fraction (44.5%, 73*/*164, one-way ANOVA, per neuron Bonferroni-adjusted p< 0.05) of neurons in the motor cortex to be tuned to paw trajectories (Fig. 5**d**,**e**,**f** and Fig. S. 7**c**,**d**). Under normal freely moving conditions, this tuning was masked by the influence of the variable body posture information, resulting in only 17.68% (29*/*164) tuned neurons. On a population level, virtual head-fixation unraveled clean neural trajectories (Fig. S. 7**e**). For some neurons (14.0%, 23*/*164), the predicted head-fixed response was similar to the actual measured response (Fig. 5**e**), potentially due to a smaller impact of body posture coding or particularly strong paw tuning in those neurons. For other neurons, no difference in the average response was detected with the classic statistical approach, whereas a trajectory tuning was unraveled via the GAM approach (27.4%, 45*/*164 neurons Fig. 5**f**, Fig. S. 7**b**,**d**).

## 3. Discussion

We adopted an approach to measure and account for movement-related information in neuronal signals. To do this, we combined 3D tracking of freely moving rats via FreiPose with a “virtual head-fixation”, which accounts for neural variation due to changes in body and head posture. Thus, we could identify a large portion of neurons in the motor cortex as being modulated by paw motions during freely moving uninstructed behavior. In contrast, the widespread multiplexing of information regarding the animals’ posture in the cortex masked this modulation when approached with classical statistical tools.

These findings were enabled by a new technical tool–marker-free holistic 3D pose reconstruction and tracking of individual body parts of freely moving animals. Rather than triangulating 2D pose reconstructions (Günel et al., 2019; Mathis and Mathis, 2019; Karashchuk et al., 2021; Nath et al., 2019), FreiPose directly reconstructs body poses in 3D, resulting in higher tracking accuracy than that achieved using previous tools. Analyzing the problem holistically by fusing information from all views into a joint 3D reconstruction allowed us to surpass the commonly used methods by 49.4% regarding the median error in freely moving rats. Marshall et al., too, argued that 2D convolutional networks are not sufficient for addressing the problem of 3D tracking across naturalistic behaviors in freely moving animals (Marshall et al., 2021). Compared to the marker-based approach, FreiPose is flexible in defining keypoints, which allows investigators to apply it to different animal models or to highly resolved individual body parts. In many such applications, markers cannot be used due to size and motion limitations. FreiPose rules out any potential effect of markers on the behavior or neural signal of the animal.

Moreover, DANNCE(Dunn et al., 2021) also showed substantial improvement versus DLC using native 3D keypoint prediction. On an abstract level, DANNCE and FreiPose employ similar strategies to elude the 2D limitations. Instead of the error-prone prediction of keypoints in individual views, both use a 3D voxel representation whereby features from individual views are projected. Subsequently, a 3D CNN reasons about the keypoint location in metric space. DANNCE constructs 3D volumes from the raw images of each view. This approach leads to a dense projection of images into the voxel space. Because the projections from multiple views overlap, an averaging out of important information can occur. In contrast, FreiPose projects sparse, learned features into voxel space (i.e., it is learned which information to best keep and project to voxel space). DANNCE used a very large marker-based ground truth dataset for rodents and had to be adapted to work on marker-less animals via retraining. In contrast, FreiPose was natively developed for marker-less tracking and, as such, does not contain any species bias. Thus, it can be adapted to novel tasks or species with a small number of hand-labeled frames (e.g., FreiPose only required 78 hand-labeled time points for the adaptation to the detailed tracking of an individual paw of a rat).

In a series of experiments, we showcased possible applications of FreiPose. In an optogenetic experiment, we demonstrated that FreiPose can control for the unintended motor effects (Aravanis et al., 2007; Gradinaru et al., 2009; Ebina et al., 2019; Sun et al., 2021) of the stimulation. FreiPose enabled the quantification of overt movements due to optogenetic perturbations in the motor cortex with minimal additional experimental effort. In a similar manner, behavioral differences between diseased and healthy individuals or effects of drugs could be analyzed and the recovery status monitored. For example, paw trajectories during reaching were used when attempting to quantify stroke recovery (Wahl et al., 2014).

We further used the tracking results for automatic clustering of behavior (Pereira et al., 2019; Dunn et al., 2021) and obtained intuitively meaningful clusters. However, the GAM approach revealed that only a small fraction of the neural variation can be explained by the behavioral state. Most of the variation was a result of differences in the exact movement during those behaviors. This finding indicates that conserved neuronal representations of behavioral categories in motor cortex (Melbaum et al., 2021) could be based on conserved neuronal representations of movements.

FreiPose also allowed us to extract single paw trajectories. We found a subset of neurons with clear tuning to the paw movement directions. To further investigate the neural responses to paw movements in the light of the observed multiplexing of different information sources in individual neurons, we presented a “virtual headfixation”, which normalizes out the body posture after the actual experiment, during which time the animal can behave naturally. With virtual head-fixation, we detected in the order of 2.5 times more paw movement tuned neurons than was detected by classical statistical approaches. These neuronal correlates were astonishingly clear and similar to those typically observed in physically head-fixed stereotyped behavioral tasks, such as center-out reaching tasks in non-human primates (Kakei et al., 1999; Moran and Schwartz, 1999; Lara et al., 2018). As such, the neural population trajectories of the “head-fixed” data, visualized via jPCA(Churchland et al., 2012), revealed a smooth directionality linked to the paw trajectories (Fig. S. 7**e**). This observation indicates that the paw encoding in a freely moving condition resembles movements under constrained conditions, thus enabling us to form a conclusion across these different experimental approaches.

The virtual head-fixation in freely moving animals extends previous attempts to describe the individual contributions of movements and other task-unrelated modalities during physically head-fixed experiments (Musall et al., 2019; Stringer et al., 2019; Salkoff et al., 2019). Our approach with behavioral characterization from FreiPose can describe the neural activity with similar precision to these previous studies but under freely moving conditions (Musall et al., 2019). This ability is remarkable because our approach works during presumably more complex and varied freely moving exploratory behavior compared to movements of head-fixed mice. Our method profits from a more precise description of the kinematics of movements, which influence the neural activity when compared to less discriminatory video motion energy or coarse motion analysis. Strikingly, the self-paced movements during exploratory behavior were represented to a similar degree to “uninstructed” movements in the cortex, as described by Musall et al. (2019). Although we showed that movements account for a considerable degree of neural variation, the source of the additional unexplained variance in those models and differences in magnitude between the GAM prediction and the observed firing rates remains elusive. We suspect that the unexplained variance is based on a mixture of three different effects: (1) neural network effects, whereby the activity of individual neurons is influenced by ensembles or connected neurons, (2) sensory processing (e.g., tactile and proprioceptive input) (Hatsopoulos and Suminski, 2011), and (3) internal motivational and planning components of non-executed movements or general behavioral states (Allen et al., 2019). In future studies, these effects can be tested by increasing the electrode count, adding more recording areas (e.g., prefrontal and sensory cortex), accounting for the potential sensory inputs from the environment.

While employing more complex methods, such as multiplicative models (Angelaki et al., 2020) or deep learning (Pandarinath et al., 2018), to analyze the neural response might provide better descriptions of the neural dynamics, they also complicate the back-projection of the predictor-response relationship onto behavioral features. The GAM approach is specifically suited to extract the contribution of individual behavioral features to neural dynamics. In our data, GAM described the neural response best during virtual head-fixation around the movement onset. The reason for this temporal effect is the focus of the model on the paw-movement variables because the paw velocity was typically highest around movement onset. To further improve the modeling, a dynamical integration of not only current movements and body postures but also of the short-term history of those variables could be employed.

Our approach serves the view that behavioral states and uninstructed movements should be taken into account during neuronal recording studies in awake behaving animals (Musall et al., 2019). Being able to conduct studies under freely moving conditions while having maintaining full control of all body parts is a large step towards avoiding the seemingly inevitable trade-off between natural conditions and scientific control. Allowing animals to move freely also has an additional positive side-effect: it simplifies training efforts because animals are often more willing to participate in complex tasks. With this approach, we revealed that robust movement representations of individual body parts are masked in the motor cortex. Because multiplexing is a widespread phenomenon in the brain (Gire et al., 2013; Ramakrishnan et al., 2017; Kremer et al., 2020), this suggests that other representations of sensory or cognitive factors might also be masked throughout the brain. The technical developments introduced here allow these questions to be addressed in a broad range of studies.

## 4. Acknowledgements

We thank the three anonymous reviewers for valuable feedback and David Eriksson for establishing the electrophysiology system that was used in this study. Additionally, we thank David Hildebrand for providing us with the marmoset dataset. This work was supported by the Baden-Wuerttemberg Stiftung, project RatTrack and the cluster of excellence BrainLinks-Brain-Tools (EXC 1086) to T.B. and I.D as well as the Bernstein Award 2012 (01GQ2301) and ERC Starting Grant OptoMotorPath to I.D.

## 5. Contributions

C.Z., A.S., T.B., and I.D. conceived and designed the study and wrote the manuscript. C.Z. implemented FreiPose and analyzed the optogenetic data. A.S. conducted electrophysiological recordings, viral injections, fiber implantation, and optogenetic stimulation, and analyzed the neural and behavioral data. M.A. implanted animals with laminar probes. F.S. recorded and labeled the mouse dataset.

## STAR Methods

### RESOURCE AVAILABILITY

#### Lead contact

Further information and requests for resources and reagents should be directed to and will be fulfilled by the lead contact, Ilka Diester.

#### Materials Availability

This study did not generate new unique reagents.

#### Data and Code Availability

The data reported in this paper (e.g., raw video data) will be shared by the lead contact upon request. Example datasets for FreiCalib and FreiPose are available via our GIN repository: https://gin.g-node.org/optophysiology/FreiPose.

Docker runtimes and source code for FreiPose will be made publicly available on GitHub: https://github.com/lmb-freiburg/FreiPose-docker. Docker runtime and source code for FreiCalib are publicly available: https://github.com/lmb-freiburg/FreiCalib. Source code for RecordTool is publicly available: https://github.com/lmb-freiburg/RecordTool. Code and examples related to the virtual head-fixation will be made publicly available on GitHub: https://github.com/Optophys/virtual_headfixation.

The remainder of the code used for analysis were made using standard approaches in python3.6 and open source code libraries, these as well as any additional information required to reanalyze the data reported in this paper are available from the corresponding author on request.

### EXPERIMENTAL MODEL AND SUBJECT DETAILS

Experiments were performed using female Long-Evans and Sprague Dawley rats aged (13-40) weeks. Rats were obtained from Charles Rivers (Strains 001,006). Female mice (20 weeks) from concurrent experiments were used solely for video recordings and obtained from CEMT, University Freiburg. All animal procedures were approved by the Regierungspräsidium Freiburg, Germany. The animals were housed in a 12 h light–dark cycle (light off from 8 a.m. to 8 p.m.) with free access to food (standard lab chow) and water. Three to four rats were pair-housed in type 4 cages (1500U, IVC type 4, Tecniplast, Hohenpeißenberg, Germany) before implantation and single-housed after the implantation in type 3 cages (1291H, IVC type 3, Tecniplast, Hohenpeißenberg, Germany).

### METHOD DETAILS

#### Motion capture

For motion capture, we combine a bounding box detection network to extract the region of interest from the full-scale images 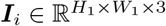 with a novel pose reconstruction architecture (Fig. 1**c**). The approach resolves ambiguities after integration of information from all views. Due to occlusion, it is typically impossible from a single view to measure the precise location of all body keypoints in that view, yet existing methods attempt to reconstruct the keypoint locations in the images, regardless of their visibility. This approach favors learning priors to hallucinate the invisible keypoints over measuring their location, subsequently diminishing performance in the subsequent 3D lifting step. To circumvent the problem, FreiPose extracts features 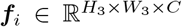 rather than keypoints from the cropped images 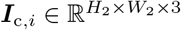 and deploys a differentiable inverse projection operation Π^−1^ (Zimmermann et al., 2019), which maps features into a 3D representation

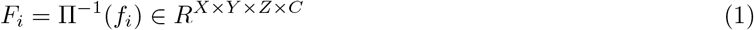

based on the features *f*_*i*_ of camera view *i*. In this notation, *H* and *W* represent spatial dimensions and *C* the number of channels for feature representation, which throughout this work are chosen as the following: *H*_2_ = *W*_2_ = 224, *H*_3_ = *W*_3_ = 28 and *C* = 128. *N* denotes the number of keypoints, which is 12 for freely roaming rodents and 14 in the reaching experiment. The input image resolution *H*_1_ and *W*_1_ lies between 600 and 1280 pixels due to varying image resolutions captured by the cameras deployed. The representations across views are merged by averaging across views ***F*** = 1*/N* ∑_*i*_(***F***_*i*_) and deploying a U-Net-like encoder-decoder architecture 3D CNN (Falk et al., 2019) on the voxelized representation. The 3D network learns to reason on the joint representation and reconstruct an initial 3D pose ***P*** ^0^ incorporating information from all views. In a similar vein to the commonly used map representation for 2D keypoints (Zimmermann and Brox, 2017), we use a voxelized representation to localize each keypoint in 3D. To obtain a point estimate, the location of the voxel with the highest score is retrieved, and we call the maximal score the predictions’ confidence *c*_*i*_.

The pose ***P*** ^0^ ∈ ℝ^*N ×*3^ is a matrix representing the location of the *N* predefined body keypoints at a given time in world coordinates. Subsequently, we use 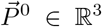 as a single keypoint sliced from ***P*** ^0^, or ℝ^4^ in its homogeneous coordinate form if required. Similarly, 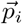 denotes a single 2D keypoint in ℝ^2^ taken from ***p***_*i*_ ∈ ℝ^*N ×*2^ of camera view *i*.

For refinement, the initial 3D pose 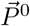 is projected into the camera views

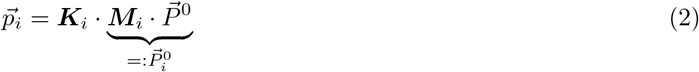

using the cameras’ intrinsic ***K***_*i*_ ∈ ℝ^3×3^ and extrinsic matrices ***M***_*i*_ ∈ ℝ^3×4^, which are obtained via the camera calibration procedure.

Given the initial 2D pose 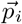 and image features ***f***_*i*_ from view *i*, subsequent convolutional layers estimate the refined 2D coordinates 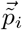. To obtain the final 3D reconstruction 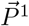, the refined 2D keypoints are unprojected into the world using:

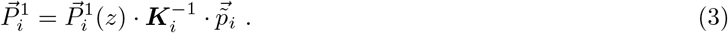

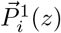 retrieves the third component from the pose in camera coordinates 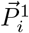, which corresponds to the respective keypoints’ depth in this camera’s coordinate frame. Secondly, the scalar reconstruction confidence *c*_*i*_ is used to calculate the final estimate as a confidence-weighted average:

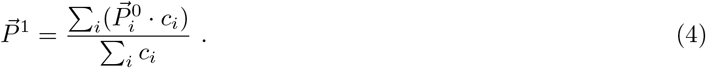

Extensive details regarding the architectural choices and algorithmic hyperparameters are mentioned when applicable throughout the Methods section (see Architectural Details of FreiPose section) or can simply be taken from the released code.

FreiPose employs a tracking-by-detection approach. Thus, each individual set of frames from all cameras is processed individually, and the temporal history of keypoints is not considered. The spatial relationships of individual keypoints are not explicitly built into the model; however, some spatial priors might be learned by the model during training.

The DLC network (based on resnet-50) was trained (for 1030000 steps) and evaluated on the same extracted bounding boxes as FreiPose. This step already strongly improved the performance of DLC compared to the base version with training and evaluation on whole images. Keypoint predictions with a confidence below *c* = 0.1 were discarded. Additionally, a RANSAC-based triangulation method was used to obtain 3D results from DLC predictions. This method was specifically developed to be less influenced by outliers. However, we would estimate that also further triangulation methods (e.g., pairwise triangulation per camera pair) would provide very similar results and only in some cases where views between some cameras disagree, RANSAC would provide better results. Furthermore, our triangulation-approach comes as a part of a full pipeline for estimation of camera parameters and calibration (FreiCalib).

#### Rat surgery

Animals were anesthetized with isoflurane (CP-Pharma, Burgdorf, Germany) inhalation. Subsequently, they were positioned in a stereotactic frame (World Precision Instruments, Sarasota, FL, USA), and their body temperature was kept at 37 °C using a rectal thermometer and a heated pad (FHC, Bowdoin, USA). Anesthesia was maintained with approximately 2 % isoflurane and 1 l/min O_2_. For pre-surgery analgesia, we subcutaneously (s.c.) administered 0.05 mg/kg buprenorphine (Selectavet Dr. Otto Fischer GmbH, Weyarn/Holzolling, Germany) and carprofene (Rimadyl, Zoetis, Berlin, Germany). Every other hour, the animals received s.c. injections of 2 ml isotonic saline. Moisturizing ointment was applied to the eyes to prevent them from drying out (Bepanthen, Bayer HealthCare, Leverkusen, Germany). The skin was disinfected with Braunol (B. Braun Melsungen AG, Melsungen, Germany) and Kodan (Schülke, Norderstedt, Germany). To perform the craniotomy, the skin on the head was opened along a 3-cm-long incision using a scalpel. The exposed bone was cleaned using a 3 % peroxide solution. Self-tapping skull screws (J.I. Morris Company, Southbridge, MA, USA) for implant stability and for extracellular referencing (above cerebellum) were placed on the skull. Craniotomies (1×1 mm) were drilled over the prospective implantation sites. For electrophysiological recordings, 7 female Sprague Dawley rats (13–40 weeks, Charles River, Germany) were implanted with 32 channel laminar probes. These animals were also involved in a different study. We implanted 1 or 2 2-shaft silicone probes (E32+R-150-S2-L6-200 NT, Atlas Neuroengineering, Leuven, Belgium) unilaterally in the motor cortex (in either RFA, CFA or both) and connected the probes via a custom interface board to allow the electrophysiological recordings via ZD32 digital headstages (Tucker-Davis-Technologies, Alachua, FL, USA). The craniotomy was sealed with Kwik-Cast (World Precision Instruments, Sarasota, FL, USA), and the implant was fixed using dental cement (Paladur, Kulzer GmbH, Hanau, Germany). The animals were given at least one week to recover.

Neural recordings
| Animal | Area | AP | ML | DV |
| --- | --- | --- | --- | --- |
| **A** | RFA | 3.35 | 1.9 | 2.4 |
|  | CFA | -0.5 | 2.3 | 3.3 |
| **B** | RFA | 3.5 | 2.5 | 2.3 |
|  | CFA | 0.87 | 2.5 | 2.26 |
| **C** | RFA | 3.27 | 2.02 | 2.7 |
|  | CFA | 0.13 | 2.43 | 2.39 |
| **D** | RFA | 3.48 | 2.07 | 2.6 |
|  | CFA | -0.6 | 2.0 | 2.5 |
| **E** | CFA | -0.5 | 2.39 | 2.4 |
| **F** | RFA | 3.5 | 2.3 | 2.4 |
| **G** | RFA | 3.5 | 2.0 | 1.67 |

Optogenetics
| Animal | Area | AP | ML | DV |
| --- | --- | --- | --- | --- |
| **1** | CFA | 0.25 | 1.9 | 1.5 |
|  | CFA | 0.45 | 2.1 | 1.5 |
| **2** | CFA | 0.2 | 2 | 1.5 |
|  | CFA | 0.65 | 2.1 | 1.5 |
| **3** | CFA | 0.3 | 1.85 | 1.5 |
|  | CFA | 0.5 | 2.1 | 1.5 |

##### Implantation and injection coordinates in mm relative to bregma

For the optogenetic perturbation experiments, 3 female Sprague Dawley rats (14 weeks, Charles River, Germany) were unilaterally injected with a ChR2-carrying viral vector (rAAV5/ hSyn-hChR2(H134R)-eYFP-WPREpA (RRID:Addgene 26973), 2.9 × 10^12^ vg/ml, UNC vector core, Chapel Hill, NC, USA) in 2 AP sites with 500 nl each using a 10 *µ*L gas-tight Hamilton syringe (World Precision Instruments, Sarasota, FL, USA). Coordinates corresponded to the caudal forelimb area (Neafsey and Sievert, 1982). Injections were performed at a speed of 100 nl/min, with a 10-min waiting period after each injection. We implanted 200 *µ*m-core fibers (NA 0.39) with metal ferrules (230 *µ*m bore size, both from Thorlabs, Newton, NJ, USA) at a depth of 1.4 mm between the injection sites. The craniotomies were covered with Kwik-Cast silicone and ferrules secured with super bond (Sun Medical Co., Moriyama City, Japan) and Paladur dental cement. Animals were allowed to recover and express ChR2 for 4 weeks before starting the stimulation.

#### Optogenetic perturbations

For the connection to the light source (Luxx 473 nm laser, Omicron Laserage, Rodgau, Germany), we employed ceramic sleeves (ADAF1, Thorlabs, Newton, NJ, USA) attached to an elastic spring-suspended custom-made patch cord with a rotatory joint (Doric Lenses Inc., Quebec, QC, Canada) to allow the rats to roam freely. Rats were stimulated for 5 sec with 30 or 10 Hz square pulses with 10 % duty cycle and 7 mW at the fiber tip with at least 45 sec between stimulation sequences.

#### Neural recordings

Broadband signals were simultaneously recorded via 1 or 2 digital head stages (ZD32, Tucker Davis Technologies, USA) connected via a flex-style dual head stage adapter (Intan Technologies LLC, Los Angeles, CA, USA) and electrical commutator (ACO64, Tucker Davis Technologies, USA) to the recording controller (Intan Technologies LLC, Los Angeles, CA, USA). Spike sorting was performed using Mountainsort (Chung et al., 2017). Spikes were synchronized to the videos via the camera frame trigger signals.

#### Video recordings

The behavioral box for freely moving exploratory behavior was made from 8-mm-thick acrylic glass and with a size of 45 × 36 × 55 cm. We used either transparent acryl or a metal mesh with 0.7 cm spacing as the floor of the box so that videos could also be recorded from below.

We used 7 color cameras (6 x acA1300-200uc and 1 x acA800-510uc, Basler AG, Ahrensburg, Germany) placed around the behavioral box to ensure visibility of the rats’ body parts from almost all angles. Ideally, at any time point, the keypoints of interest should be visible from multiple cameras to ensure accuracy (for more details, see GitHub tutorial). We employed objectives with a focal length of 6–8 mm (Kowa, Nagoya, Japan). To calibrate the cameras, we employed a calibration pattern (described in FreiPose framework in Supplementary). To ensure a planar surface and exact marker size, the pattern was manufactured by Schmidt Digitaldruck GmbH, Wörth, Germany.

The cameras recorded with 30 fps using custom python software. The software is able to set exposure time, gain, and frame rate, and can adjust the white balance. It also allows for individual camera control and recording of calibration sequences. However, FreiPose would work with videos recorded with any other recording systems as long as synchronous frame acquisition is ensured. We used a low-cost Arduino-based hardware trigger (details and Arduino code available on GitHub) for frame synchronization. The frame trigger signals were also recorded with the Intan recording system to allow for synchronization of the video with neural signals.

##### Tracking of paw during reaching

We trained a rat in a reaching task. The animal was food-deprived (limited to 3 standard pellets) prior to reaching training. It was then placed in a transparent box (27 × 36 × 29 cm) with an opening (1 × 6 cm), in front of which sugar pellets were placed. For the reaching task, we used a lens system with a larger focal length (12–16 mm, Computar, Cary, USA / Cosmicar, Tokyo, Japan) to focus the camera’s image resolution on the reaching movement.

##### Tracking of mice

We trained FreiPose to predict keypoints on mice. C57BL/6 mice, implanted with a miniscope (from concurrent experiments in our lab) were placed in a transparent box made from 5mm acrylic glass (22 × 13.5 × 25 cm), which they explored. To ensure visibility of all body parts, four monochromatic cameras (4x Basler a2A1920-160umBAS, Basler AG, Ahrensburg, Germany) were placed around the behavioral setup, one below and three from the sides. We employed objectives with a focal length of 4–8 mm (Basler AG, Ahrensburg, Germany). The cameras were recorded with 30 fps using PylonViewer (Basler) and the Arduino-based hardware trigger described above. To enable synchronization of neural signals with camera recordings, the frame trigger signals were recorded using an OpenEphys Acquisition box.

#### Architectural Details of FreiPose

Our approach was implemented using Tensorflow (Abadi et al., 2016) and is available through our GitHub repository.

For bounding box detection, we used a COCO (Lin et al., 2014) pretrained MobileNet V2 (Howard et al., 2017), which was retrained for the task of detecting the foreground objects. In the freely moving rat scenario, it was trained using each view of the 1199 labeled time instances separately (i.e., a total of 9592 samples). We trained it for 150 k iterations using a learning rate of 0.004 and the RMSProp optimizer. As data augmentation operations, we employed random flipping, cropping, scaling, and color space variation. For the reaching task, because the action always occurred in the same space, we used a fixed, predefined region of interest rather than a detector.

For pose estimation, we extend the approach by Zimmermann et al. (2019) using the refinement module for increased accuracy. The network was trained for 60 k using ADAM optimizer (Kingma and Ba, 2014) with a base learning rate of 10^−4^ and decay by a factor of 0.1 every 30 k steps. To improve convergence, we found it helpful to not train the refinement module for the first 30 k steps. Furthermore, we used a 64 × 64 × 64 voxel grid for 3D volumes.

#### Skeletal models

During the freely moving rat experiment, we use a 12 keypoint model (Fig. S. 8**a**), which includes keypoints along the body axes, faces, and paws (Fig. S. 8**b**). During the freely moving mouse experiment, we utilized a 10 keypoint model, similar to that used for the rat experiment, excluding the eyes. For the reaching experiment, we defined a 14 keypoint model using a point at the wrist, one for the thumb and three for all the remaining fingers (at metacarpophalangeal, proximal interphalangeal joints, and fingertip). For the bird experiment, we used a 12 keypoint model as defined by the authors of the dataset (Badger et al., 2020). For the marmoset experiment, we employed a 18 keypoint model as defined by the authors of the dataset (Dunn et al., 2021).

#### Details of generalization experiments

For the reaching experiment, we labeled 108 frames and used 78 as training and 30 as evaluation. The avian dataset(Badger et al., 2020) was obtained from https://marcbadger.github.io/avian-mesh/, while the marmoset dataset was kindly provided by David Hildebrand (Dunn et al., 2021). For both datasets, the calibration data, as well as the labeled keypoints, were transferred into FreiPose notation. Some labels were further adapted due to an insufficient quality of single samples. The cowbird dataset consisted of 144 labeled frames from 8 cameras and was subdivided into 126 training samples and 18 evaluation frames. Marmoset images contained 92 labeled frames from 3 cameras and were split into 72 training samples and 20 evaluation frames. Since this dataset was also used in Dunn et al. (2021), the interested reader can compare our Fig. 2**f** with the figure panel 6d from Dunn et al. (2021) for a quantitative comparison. Please note that different train/test splits were used in the current manuscript compared to DANNCE. For the mouse, we labeled 319 frames from 4 cameras. For all models, we used three random folds to define the train/evaluation split for network evaluation.

For the species generalization experiments, FreiPose was trained for 80 k (100k for mice) using ADAM optimizer (Kingma and Ba, 2014) with a base learning rate of 10^−4^ and decay by a factor of 0.1 every 30 k steps. For the base rat net, we used a voxel resolution of 0.4, for mice 0.1, and 0.8 for birds and marmosets. Voxel resolution is dependent on the size of the object to be tracked as well as the total volume where the object can be found. Because the voxel resolution is limited, the larger the animal’s enclosure, the lower the maximally achieved accuracy.

For the avian model, no bounding box network was trained due to the presence of multiple animals. A modified bounding box network for the detection of individual animals could be envisioned in the future and seamlessly implemented with FreiPose.

### QUANTIFICATION AND STATISTICAL ANALYSIS

#### Quality metrics

Building on the notation introduced in Motion capture section, the median error is calculated as follows: Let 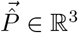 denote the predicted keypoint coordinate of one keypoint and 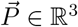 represent the label; then, the reported metric is defined as

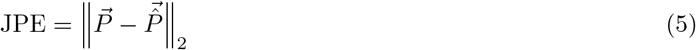

and represents the **J**oint **P**osition **E**rror (JPE); JPE were calculated over all trials, evaluation frames, and keypoints.

The percentage of correct keypoints (PCK) was calculated using different thresholds assessing the fraction of keypoints with an error below the corresponding threshold.

#### 1D tuning curves and postural variables

Tuning curves were based on free exploration behavior. Due to the lack of a clearly defined point on the rat’s fur corresponding to its skeletal neck, we used the midpoint between the ears as the surrogate neck. We defined the body axis of the rat as the vector from the tail root to the midpoint between the ears, and the head axis was demarcated as the vector from the midpoint between the ears to the nose. Body and head pitch were calculated as the angle between the horizontal and the body or head axis, respectively. Angles between resulting vectors were calculated using trigonometric functions.

To calculate the tuning curves, the head pitch and azimuth were binned in 4 degree steps, body pitch in 2 degree steps. Neural firing rates were binned according to the bin edges (in time) of the body pose variables. By shifting the neural data relative to behavior by ± [5 : 60] s 1000 times, we obtained a shuffled population. Neurons were considered significantly tuned when their extrema exceeded a z-score corresponding to a Bonferroni-adjusted p-value of 0.05 (quantile function of normal distribution at (1 − 0.05)*/N*) of the shuffled population for at least 3 consecutive postural bins.

For some postural variables (e.g., paw movements), it was necessary to transform the location of the keypoints to the rat’s local reference frame (such that the rat’s body axis was oriented along the calculated x axis). This approach distinguishes the movements of the paw relative to the body from the absolute movements of the paws while the entire body is moving (e.g., during rearing). A rotation matrix was created based on the body axis and vertical axis (pointing up). Subsequently, keypoints were multiplied with the rotation matrix to obtain keypoints in the new reference frame.

To express movements or postural directions, the direction vector is projected onto a plane (e.g., for the xy-direction of a paw movement, by removing the z-component for the horizontal plane projection). The angle to the ordinate was calculated via trigonometrical functions; the histogram of the resulting angles was smoothed with a Gaussian filter for representation purposes and plotted onto a polar projection.

#### Supervised behavioral classification

To subdivide the continuous sequence of behaviors occurring during free exploration, we developed a supervised machine-learning approach. To this end, we annotated 5 sessions from different animals through visual inspection using MuViLab (https://github.com/ale152/muvilab). The video from a single camera was divided into 1 s snippets, and each snippet was categorized as either locomotion, grooming, exploration, rearing, sitting, or unclear. Locomotion was defined as longer, typically straight movements. Time points with shorter movements in a stop-and-go fashion, as well as direction changes, were defined as exploration because, typically, this exploration was accompanied by sniffing and whisking. Grooming clusters were characterized by a typical sequence of paw-licking, nose and ears cleaning, and fur cleaning. Rearing was defined as time points with both front paws in the air and elevated body axis, and sitting (resting) as time points with no motion or only minimal head movement. A more detailed separation of behaviors into further subclasses is possible but is beyond the scope of this work.

We then performed behavioral decomposition of the tracking data of the corresponding videos. The 3D keypoint predictions from FreiPose were transformed into 912 variables considering the position of the keypoints relative to each other as well as the frequency of their modulation (for a full table, see quantification of optogenetic stimulation). These variables were then non-linearly embedded using UMAP (McInnes et al., 2020) and reduced to 5 dimensions. Subsequently, a SVM classifier (*sklearn* library, C=1, linear kernel, with balanced class weights) was trained on the points in the embedded space (each point in the embedding corresponds to 912 variables for each frame) using the labels from the annotations from 3 training sessions, and evaluated on 2 test sessions. “Unclear” labels were omitted for training. To predict behavior in novel sessions, the same trained UMAP and SVM models were employed in order to transform the keypoint predictions.

For visual inspection of the predictions, video snippets of 1 s duration were labeled with the corresponding prediction (only if *>* 70% of the frames in a snippet were predicted to have the same label; otherwise, an unclear label was given) and viewed in MuViLab.

#### Unsupervised behavioral embedding

We modified an algorithm for behavioral mapping of freely moving fruit flies (Berman et al., 2014) to cluster the pose reconstruction time-series into behavioral clusters in an unsupervised manner. Whereas the original algorithm relied on postural decomposition via principal component extraction from videos, we employed the FreiPose-predicted body keypoints as postural information. Predicted poses were smoothed with a Gaussian (std of 33 ms, 1 time bin) and transformed into the rat body-centric reference frame. Pairwise distances of all keypoints, as well as the distance between each keypoint and the floor plane, were combined, and the top 20 principal components (explaining 95 % of variance) were extracted. Time courses of the components were Morlet-wavelet transformed with the *PyWavelets* library at the following frequencies: 0.5 and 1–15 Hz in 1 Hz steps. We then recombined the temporal features (results of the wavelet transform for all frequencies and components) with the spatial features (keypoint location in the rat’s local reference frame). Combined features were z-scored and embedded into 2 dimensions via the t-SNE approach (Maaten and Hinton, 2008). We employed the *openTSNE* python library with cosine distances and PCA initialization, perplexity 100, and exaggeration 12 as parameters. Subsequently, the tSNE embedding was smoothed via kernel density estimation with a 501 × 501 dot canvas size and a bandwidth of 0.08. Areas around peak densities were separated via the watershed (*scikit-image* library) algorithm.

To evaluate whether the behavioral clustering resulted in meaningful separations of behavior, time points dwelling in a single cluster for longer than 0.1 s were extracted from the corresponding video and concatenated into a single video file for evaluation by a human observer using *ffmpeg* (2.8.15) libraries. Because t-SNE is a non-parametric, stochastic approach, the results between runs might vary.

#### Neural firing during behaviors

Importantly, the behaviors we chose for the supervised classification algorithm are only meaningful if a short time sequence is considered because a specific movement integration has to occur. As such, we chose 1 s window as a meaningful resolution of behavior. Because we intended to analyze the neural modulation of behavior as a state and not as transitions between different behaviors, we consider this time resolution to be sufficient. A finer time resolution can be obtained using different behavioral classes or unsupervised classification methods but is beyond the scope of the current work.

To analyze whether neurons modulated their firing rate during different behavioral states, we subdivided the neural data into 1 s bins after convolution with a 100 ms Gaussian window. Only bins with stable behavior were considered. The mean firing rates during these bins were compared via one-way ANOVA according to the behavioral label (from the supervised classifier) of the corresponding bin. The p-values were Bonferroni-adjusted according to the number of neurons tested.

#### Quantifying the effect of optogenetic stimulation

To systematically investigate the effect of optogenetic stimulation in freely moving rats, we recorded the movements of three animals across four sessions—two sessions with 30 Hz laser burst frequency and two sessions with 10 Hz with a stimulation duration of 5 sec, 10% duty cycle. During each recording session, we stimulated each animal 5 to 7 times, with a minimum inter-stimulus time interval of 45 sec. We retrained FreiPose based on 136 samples from these recordings and systematically defined 908 behavioral variables for every time step from the reconstructed poses. Behavioral variables included transformation of the pose into a rat-aligned Cartesian coordinate frame, the distance between each keypoint and the ground floor, as well as their velocities and Fourier transforms. Additionally, we calculated the angles between body limbs with respect to each other and the direction of gravity (e.g., angle between head and body axis; see table for the full list of variables).

To reveal changes in behavior, we followed an *attribution-by-classification* paradigm: given the behavioral variables at a time step *t*, we trained a linear SVM model (*C* = 0.0025) to classify every time step into stimulated (i. e., *positive*) or not stimulated (i.e., *negative*). We trained separate classifiers for each animal, used one recording for training, and retained one for evaluation.

To attribute the effect of stimulation to individual body parts, we trained classifiers on a single variable level. A separate classifier was trained for each animal, burst frequency, and behavioral variable (Fig. S. 4 **b**). The significance of each feature was assessed (p-value < 0.001, Chi2 test). However, the resulting features were not easily comprehensive (e.g., “Paw Right Front: Height over Ground f=10.3Hz”). In order to visualize the results more comprehensively, we developed the lineage tracing approach, whereby the transformation path of each feature is back-traced to the originating body part. Each path on the map is weighted depending on how many significant features pass along it. The resulting tree is easily visualized and enables one to infer which body part was affected by the stimulation.

| Name | Number of factors |
| --- | --- |
| Keypoints in cartesian rat local frame | $12 \cdot 3 = 36$ |
| Keypoints' velocity in cartesian rat local frame | $12 \cdot 3 = 36$ |
| Keypoints' distance to rat local frame origin | 12 |
| Keypoints' distance velocity to rat local frame origin | 12 |
| STFT of Keypoint distance to rat local frame origin using 33 frequencies | $12 \cdot 33 = 396$ |
| Keypoint distance to ground plane | 12 |
| STFT of Keypoint distance to ground plane using 33 frequencies | $12 \cdot 33 = 396$ |
| Body elevation: $\angle(g, [11, \{0, 5\}])$ | 1 |
| Head elevation: $\angle(g, [10, \{0, 5\}])$ | 1 |
| Head roll: $\angle(g, [0, 5])$ | 1 |
| Head nick: $\angle([11, \{0, 5\}], [\{0, 5\}, 2])$ | 1 |
| Paw right: $\angle([11, \{0, 5\}], [\{0, 5\}, 2])$ | 1 |
| Paw left: $\angle([11, \{0, 5\}], [\{0, 5\}, 7])$ | 1 |
| Angle velocity of the aforementioned angles | 6 |
| Total | 912 |

**Tested behavioral variables.**∢(.) measures the angle between two 3D vectors, and [*a, b*] defines a vector that goes from point *a* to point *b*. The notation {*a, b*} calculates the average of the points a and b. 3D points are denoted as keypoint indices that find their textual counterpart in Table S. 8. The rat local coordinate frame is defined with its origin at {0, 5}, its z axis being aligned with the up pointing normal of the ground plane and its y axis rotated towards [{0, 5}, 11]. STFT–short-time Fourier transform.

#### Paw trajectory stratification

To stratify the front paw movements during freely moving behavior, we extracted periods of high velocity in relation to the body, preceded by periods of lower velocity of the paw defined by a velocity threshold. The threshold (0.012 m/frame) was chosen manually using co-inspection of the velocity trace and the animal behavior. We further tested the influence of the exact threshold by varying it by a factor log_2_[−1.5 to 1.5]. The difference in the resulting tuning was assessed via one-way ANOVA with the threshold as main effect and sum of spikes in ± [333 ms] window around movement onset as the dependent variable.

By transferring paw positions into a rat-body-centered reference frame (see above), we extracted paw movements relative to the body. 3D trajectories of the paws were smoothed with a Gaussian (std of 100 ms) and a window ranging from −66 to +500 ms around the time point of the threshold crossing that was extracted. Trajectories were aligned to begin at the origin (0,0,0) of the reference frame. We subdivided the directions of the paw trajectories via k-means clustering (k = 10, using elbow approximation) and visually analyzed the trajectories in each cluster. To obtain a clearer relationship for directionality, we chose clusters with clear horizontal components in trajectories and consistency in trajectory length and direction. Five clusters did not fulfill these criteria (e.g., with main components pointing upward or downward) and were therefore excluded from further analyses. A description of all clusters can be found in Fig. S. 5 **d**. The duration of each movement was defined as the time from the start of the movement until the time point when the velocity falls to 33% of its peak.

We stratified the neural responses according to the calculated cluster labels from the corresponding trajectory. For the raster plots, the neuronal spike times were aligned to the frame triggers (and, thus, to the behavior) and segmented into 1.1 ms bins. To obtain continuous firing rates, the spike trains were smoothed with a Gaussian (std of 33 ms).

To identify neurons that were significantly modulated to different trajectories, we calculated a one-way ANOVA per neuron (dependent variable—sum of spikes in ± [333 ms] window around movement onset, independent variables—trajectory cluster labels) and Bonferroni-adjusted the resulting p-values to account for the number of tested neurons. Neurons were considered significantly modulated if the adjusted p-value for the main effect was < 0.05.

To analyze the population response to the paw movements, the neural responses around these paw movements (± 15 bins) were extracted. A pre-trial mean and std were calculated from [-20:-10] bins relative to movement onset, along trials, and then a mean of those values across bins. Then, a z-score was calculated per bin as (*bin value* − *pre mean*)*/pre std* for each neuron. Only trajectory classes shown in the figures were used for the analysis.

#### Generalized additive models

To estimate the cumulative influence of multiple variables onto the firing rate of a neuron, we employed generalized additive models (GAM). For model fitting, a python library *pyGAM* (Servén and Brummitt, 2018) was used. GAM provides the flexibility of modeling non-linear relationships by modeling the response-predictor relationship using a combination of multiple smooth splines, yet keeping the tractability of the effects due to the linear combination of terms. Each neuron was modeled individually; the firing rate was the response variable, and a set of predictor variables was available to the model. We predefined a set of variables based on FreiPose results as well as measured neural activity. The full list of variables and their descriptions can be found in subsequent section. We used the LinearGAM model with 10 splines and a 0.6 lambda penalty for each term (variable). To account for potential differences in timing in predominantly sensory or planing neurons, we fitted multiple time-shifted models for each neuron. For each of the time-shift models, the neural data were shifted relative to the behavioral data by ± [0:2:10] bins. The model with the highest explained variance was used for further analysis.

#### Variables extraction for GAM

To provide the GAM with available information regarding ongoing behavioral and neural activity, we designed a set of variables, as well as their derivatives, based on measured neural activity and keypoint tracking by FreiPose. We defined each animal’s body and axis as described previously. Global variables and their derivatives describe the axis directions in global reference frame *head_dir_global, body_dir_global, global_velocity*, whereas the body posture variables describe the position of bodily axes in rats’ egocentrical (centered on its body axis) perspective *body_pitch, head_pitch, head_azimuth, head_angle, head_roll*. For paw variables, the movement vectors in each Euclidean axis (in egocentrical reference frame) were extracted for each paw *dXr, dYr, dZr, dXl, dYl, dZl, dXrh, dYrh, dZrh, dXlh, dYlh,dZlh*. For each of these variables, the derivative was calculated and zeropadded—*d variable*. A behavioral class prediction obtained from our classifier (see behavioral classification) was integer encoded *behaviors*.

##### Forward-search procedure

We intended to build a model of neural activity with high explanatory power. pyGAM and similar packages provide statistical estimation of the significant contribution of a given input regressor to the model. However, a statistically significant contribution does not necessarily mean a relevant contribution. Thus we decided to use the more conservative approach of forward-search and only added regressors to the model if they contributed relevantly (increase in explained variance *>* 0.05).

To choose the most appropriate variables to model a certain neuron’s response, we employed a step-wise forward-search procedure. As a first step, a model was fitted using all variables one by one. The variables were then ranked according to the explained variance of the corresponding model. The variable with the highest explained variance was kept and the next best variable added so that a new model could be calculated. This approach continued until the addition of the next variable increased the explained variance by less than 0.05; then, the forward search was stopped, and the last added variable was discarded. A toy example can be found in Fig. S. 10.

##### Isolating body posture influence

To compare the body posture tuning estimated by the GAM approach to the bootstrap approach, the response of the individual neuron was predicted over a grid of corresponding body posture variables. An individual variable was set to an evenly spaced vector within the occurred range (e.g., 0 to 45 degrees for body pitch), whereas all other predictor variables were fixed to limit their contribution. A union of the set of neurons, which were identified as coding for a certain body posture variable by the bootstrap approach, was then calculated with the set of neurons identified as having a significant modulation to the same body posture variable with the GAM approach.

To compare the obtained tuning curves using the GAM approach with the previously used bootstrapping strategy, we analyzed the union of neurons detected as tuned via both approaches. We further analyzed the resulting tuning curves on the population level for each neuron detected as tuned for both approaches. To this end, the results from both methods need to be evaluated at the same values of postural variables. We approximated and interpolated (using *scipy*.*interpolate*.*inter1d*) the resulting tuning curves over the entire observed range for all postural variables. Such resampled tuning curves could then directly be compared via correlation (*numpy*.*corrcoef*).

##### Cross-validation

Here we employed GAM as a descriptive model, whose intention is limited to the description of the data of each individual neuron/session. GAM is only used to describe the relationships of measured regressor variables (e.g., body pitch) to the firing rate of this specific neuron. To make sure no spurious relationships are picked up, we additionally checked the performance of cross-validated (cv) models. We split the data in training and test splits 10 times (ten-fold cross-validation using *sklearn*.*model_selection*.*groupkfold*). Groups were chosen as 2 s continuous pieces across the session, to ensure coverage of the whole session. The average autocorrelation of the neural signal at 2 s lag was only 0.09, ensuring sufficently different data in each group. As was expected due to strong regularization of GAMS and limited degrees of freedom, the cv-models reached very similar performance (0.144±0.097 (std) for full-data models vs. 0.110±0.092 for cv-models, see Fig. S. 6 **d**). More importantly, for the virtual head-fixation estimations, the learned regressor-response relationships are much more relevant than the exact accuracy of the model. We analyzed the regressor-response relationships between the full-data models and the cv-models and found them to be highly similar (average correlations 0.97±0.04 std). See Fig. S. 6 **e** for an example and **f** for a summary of correlations for all neurons. This confirmed that the estimated relationships between the individual regressors (e.g., body pitch) and the response variable (neural firing rate) are indeed reasonable in the full-data model. We reported the model performance as cross-validated cvR^2^.

#### ΔR^2^ contribution

To estimate the importance of individual variables to the explanatory power of the individual GAMs, we analyzed the reduction of explained variance (Δ*R*^2^) upon temporal shuffling of one or multiple variables included in the model. GAM was fitted as described previously. Using the data for fitting, we then randomly shuffled the temporal pattern of variables belonging to a particular group (e.g., front-paw right) and re-evaluated the model using these new data. Δ*R*^2^ was calculated as the difference to the explained variance before shuffling. To calculate the *R*^2^, the predictions were normalized to their mean to adjust for a potential shift in the baseline (bias). The relative group contribution was calculated by dividing the Δ*R*^2^ by the *R*^2^ of the entire model. Variables were grouped as head: [‘*head angle*’, ‘*head roll*’, ‘*head pitch*’, ‘*head azimuth*’ and their derivates], body: [‘*body pitch*’, ‘*dbody pitch*’], FrontRPaw: [‘*dXr*’, ‘*dY r*’, ‘*dZr*’, ‘*ddXr*’, ‘*ddY r*’, ‘*ddZr*’], FrontLPaw, HindRPaw and HindLPaw accordingly. Paws: all paws combined, All motion: all previous variables combined.

#### Virtual head-fixation

For the virtual head-fixation approach, a GAM was fitted for each neuron using the previously described approach, this time forcing the model to contain the paw-related variables (dX, dX, dZ, etc.) by bypassing the forward search. This strategy enabled us to estimate the neuronal response to the paw movements. During the forward search, for each model, a set of variables were extracted from the measured data, which were identified to be predictive of these neurons’ firing rate. Body posture variables were set to their median, thereby eliminating variations in body posture and providing “virtual head-fixation”. The paw variables were kept as they originally occurred. The “head-fixed” firing rate of the neuron was then predicted based on this modified dataset. This prediction constituted the reduction of the neuronal response to the paw movements.

##### Analyzing the “head-fixed” prediction

Predicted peri-movement “head-fixed” firing rates were extracted according to the movement speed threshold in the same manner as described earlier. To compare how similar and at what time points the head-fixed model best described the firing rate of neurons, we calculated the correlation (scipy.spatial.distance.correlation) between the predicted head-fixed firing rate and actual firing rate. Additionally, we determined the correlation of the postulated head-fixed behavior (behavior with body posture variation removed) to the actual behavior occurring during the same time epochs. For individual neurons, a subset of paw movement trials with high (*>* 0.5) behavioral correlation was compared to all paw movements as well as the GAM prediction for this subset. Such trials with high behavioral correlation contain behaviors that are similar to the postulated “head-fixed” posture. The neural correlations were compared via a Wilcoxon signed-rank test.

#### jPC Analysis

To visualize the neural trajectories, we employed the jPCA technique(Churchland et al., 2012). Python implementation of jPCA was used (https://github.com/bantin/jPCA). Neurons that achieved explained variance *>* 0.35 in the full model were included in the analysis. We individually fitted and projected the neural firing rate and “head-fixed” prediction during the trials onto the first two components of jPCA, which was fitted using default parameters using 1 s window centered around the movement initiation (see previous).

#### Supplemental Items

SuppData_OptoExperiment.xlsx - Supplemental Table including full results of the optogenetic quantification experiment. Related to Figure3.

## SUPPLEMENTAL ITEMS

**Figure S. 1:**
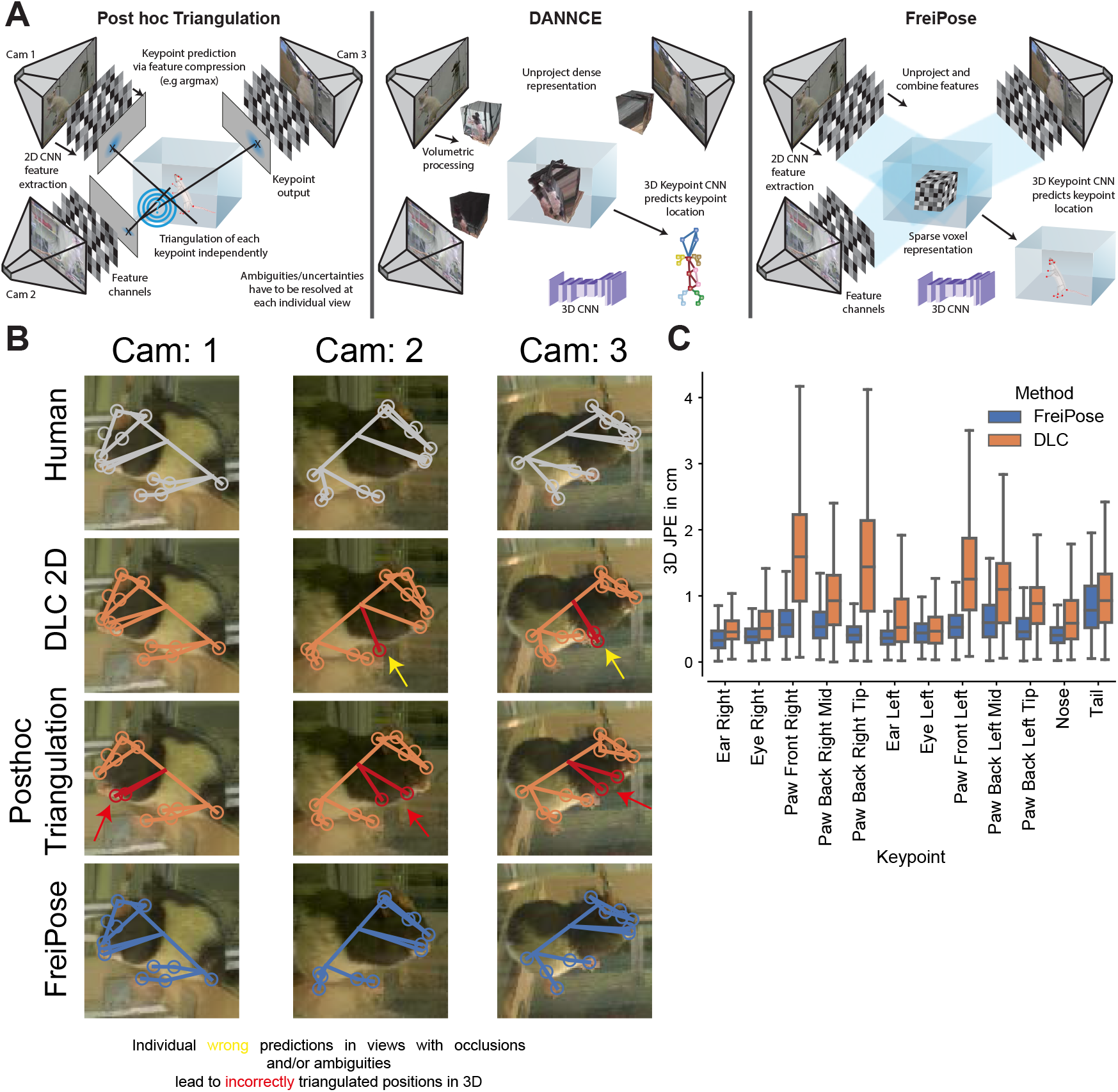
Comparison between FreiPose, DANNCE, and DLC. Related to Figure 1. Comparison of the approaches on conceptual and qualitative basis. **a** Conceptual comparison. Commonly used motion capture approaches (e.g., DeepLabCut) reconstruct 2D poses for each view independently and subsequently calculate a 3D pose (left). This post hoc triangulation requires the resolution of ambiguities in each view separately, and errors are carried over to the 3D triangulation. DANNCE (Dunn et al., 2021) uses a volumetric approach in which the dense image representation is unprojected into a common 3D space (middle). FreiPose accumulates evidence from all views (via projection of features into common 3D space), leading to a holistic 3D pose reconstruction (right). Ambiguities are resolved after information from all views becomes available. Parts of the figure were used in Fig. 1**b. b** Qualitative comparison. The DLC-based single view estimation plus post hoc triangulation was prone to erroneous predictions from individual views. DLC was run with a RANSAC-based triangulation method and confidence score cutoff to take outlier measurements into account. The triangulation method is part of the released code within the FreiPose Github repository. Despite these modifications, DLC’s predictions were not reliable. Shown are camera frames recorded at the same time but from different views. DLC (orange) was able to correctly estimate keypoints in the Cam1 view, but, during triangulation from 2D predictions to 3D points, the less accurate predictions from the Cam2/3 views had a deteriorating influence on the final results, even though robust triangulation techniques were used. FreiPoses’ (blue) predictions had a striking similarity to the annotations made by a human annotator (grey). **c** Box plot of the keypoint prediction error per keypoint for FreiPose (blue) and DLC (orange), where whiskers indicate 1.5 × IQR. The largest improvements between FreiPose and DLC were observed for highly articulated keypoints, e.g., paws.

**Figure S. 2:**
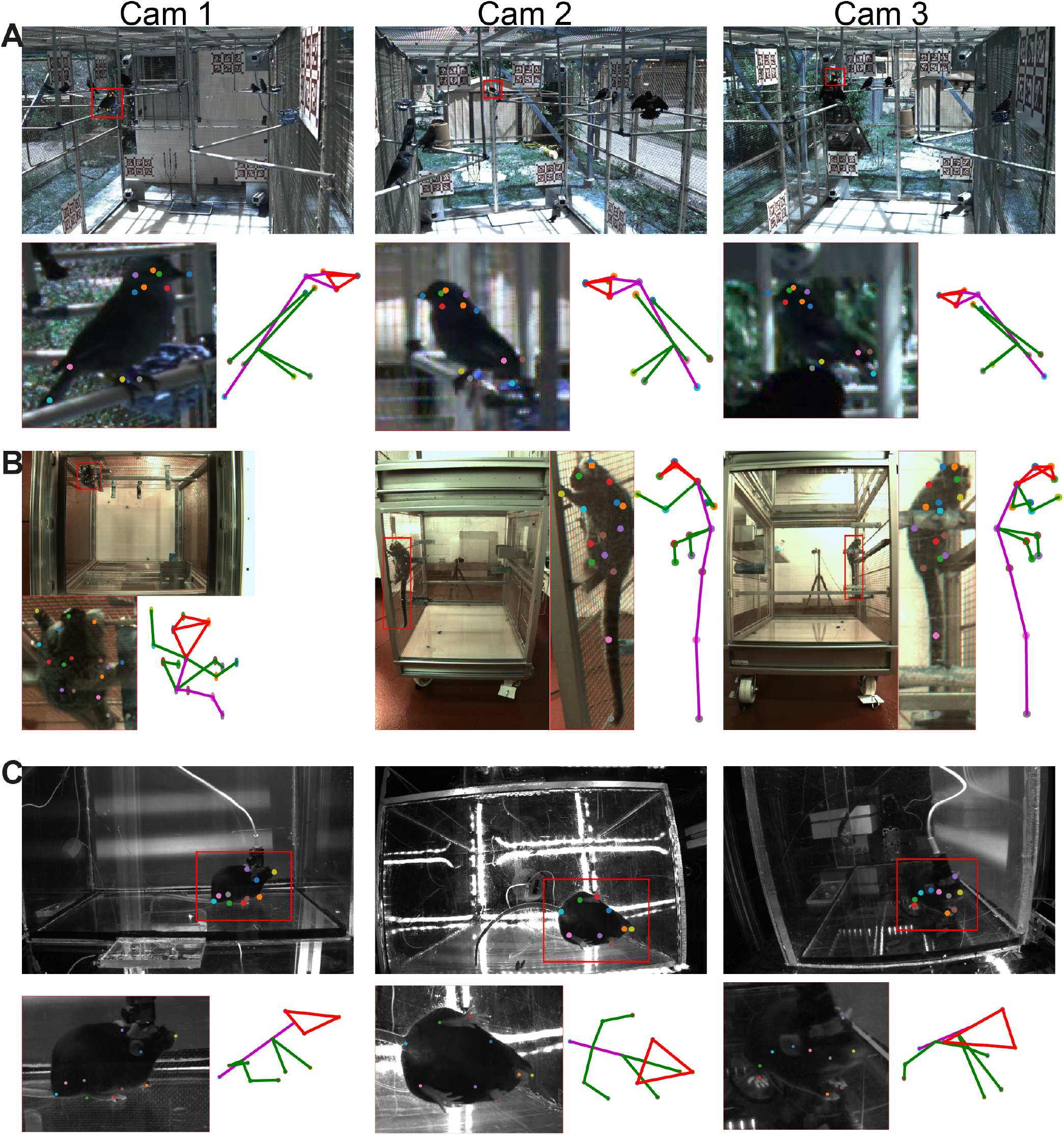
Generalization of the FreiPose for different tasks or species. Related to Figure 2. Detailed example FreiPose predictions on different datasets. Shown are individual camera views and zoomed-in views, as well as a skeletal representation of predicted 3D keypoints for 3 different camera views. **a**- Cowbird, **b**- Marmoset and **c**- Mouse.

**Figure S. 3:**
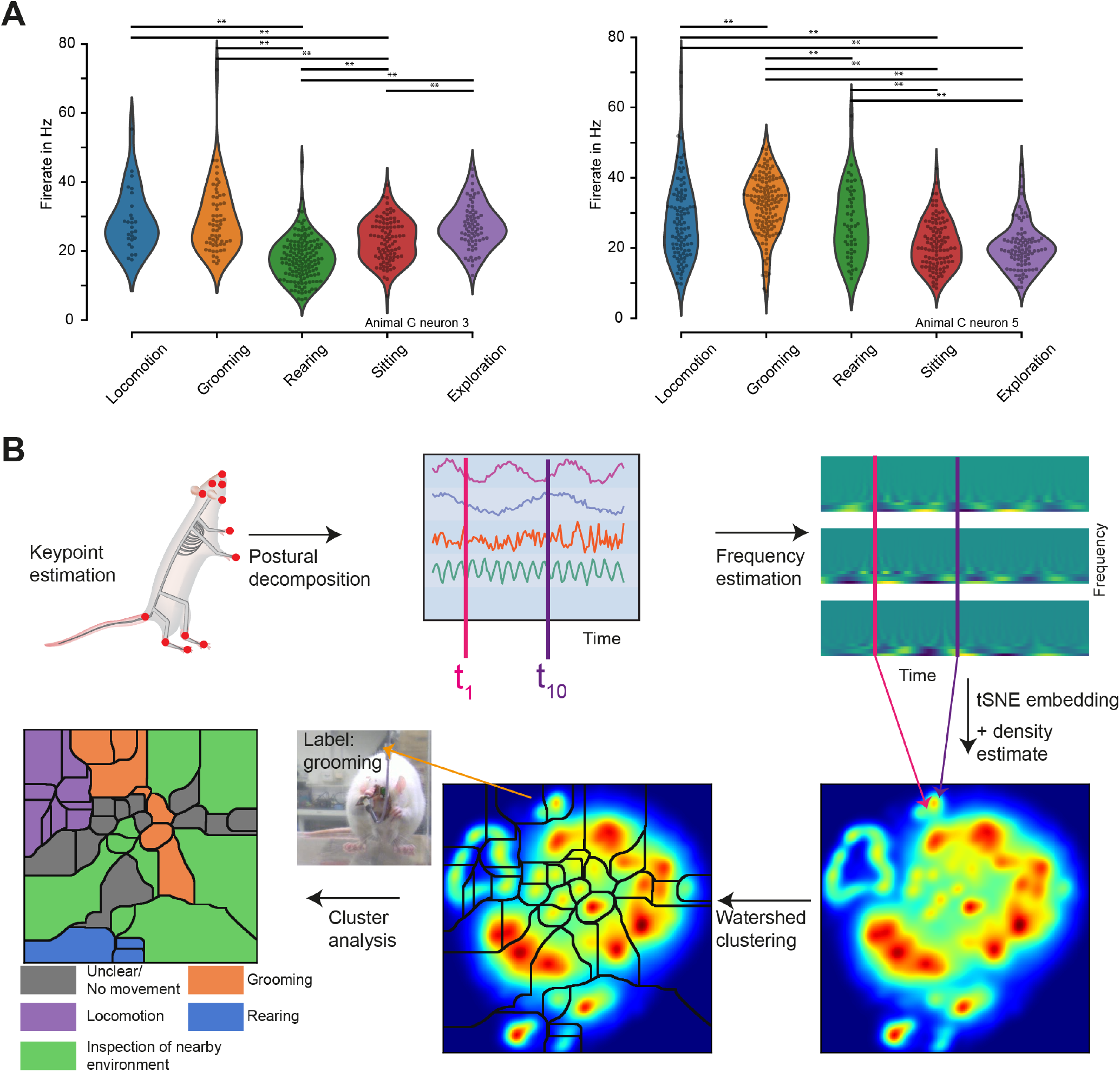
(Un)supervised clustering of behavior. Related to Figure 3. **a** Violin plots of the representative example neurons with a distinct firing rate for each behavior (one-way ANOVA per neuron with post hoc t-tests, Bonferroni-adjusted, ** indicate p < 0.01). The neuronal firing rate was evaluated for different behaviors as predicted by the supervised algorithm. The firing rate was averaged in 1 s intervals with the stationary behavioral class. **b** Algorithm for unsupervised clustering of behavior. Postural decomposition was achieved via PCA of the pairwise-distance matrix of keypoints. The frequency of the components was calculated via a wavelet transform. Extracted features were embedded via t-SNE and individual behavioral clusters identified with watershed clustering followed by assignment of clusters to behavioral labels by visual inspection and evaluation. Density estimate of the non-linear embedding and watershed clustering of a single session and cluster identities based on visual inspection of the behavior.

**Figure S. 4:**
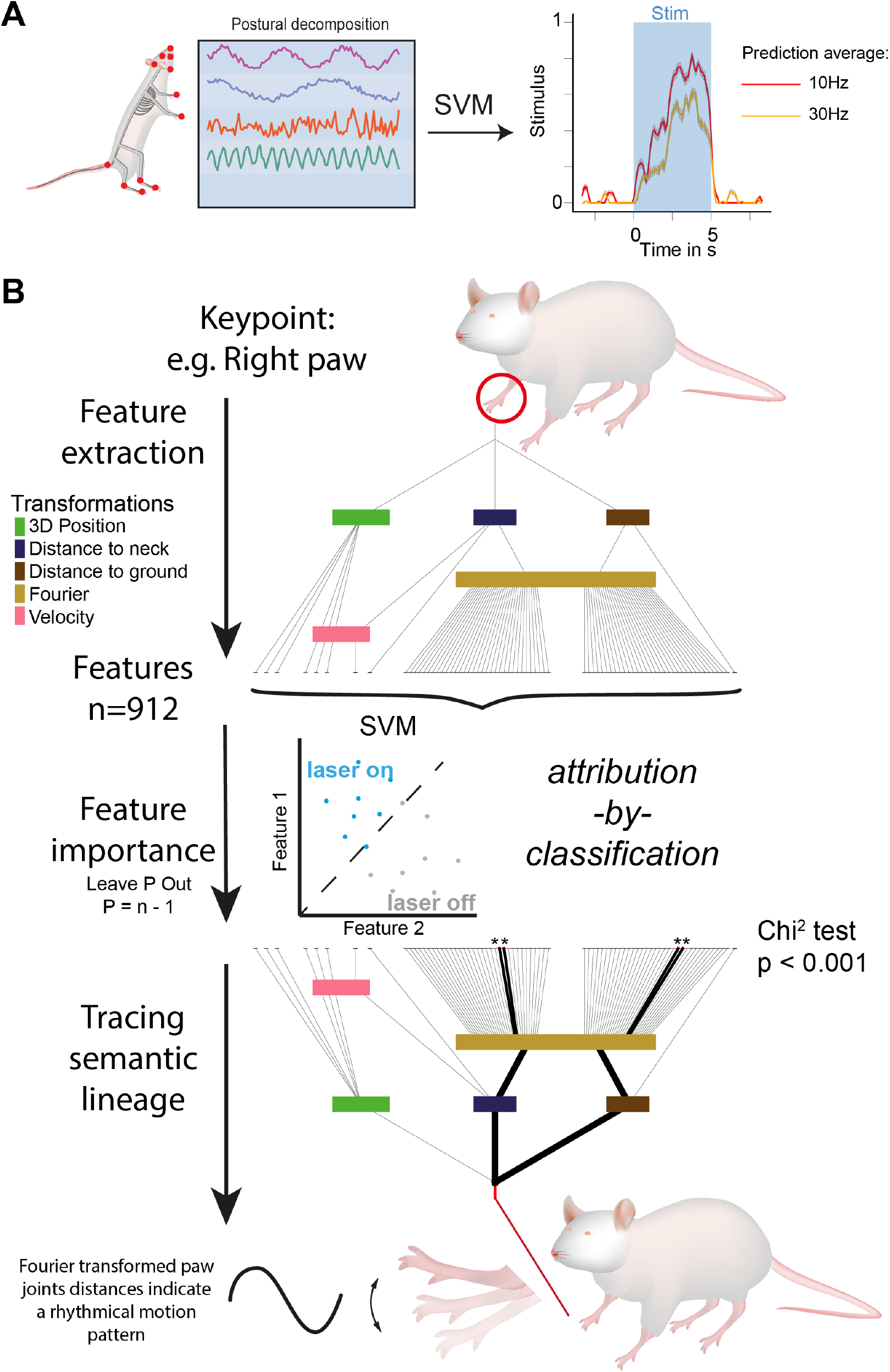
Identification of optogenetic effects using FreiPose. Related to Figure 3d. Rats expressing ChR2 in the CFA were stimulated with 5-s-long pulse trains. To detect the optogenetic effects, the pose of the animals was estimated using FreiPose and decomposed into individual features (see Methods). **a** A SVM classifier predicted whether or not a time step was stimulated (“1” or “0”, respectively). We averaged the predicted stimulus of the classifier model across stimulation trials on recordings that were withheld for evaluation. Curves were smoothed over a window length of 23 ms. Shaded areas indicate SEM and blue shaded area indicates optogenetic stimulation. **b** Lineage tracing approach for attribution of optogenetic effect to specific body parts. Decomposition of keypoint position into features involved multiple transformations described in Methods. Classifiers (SVM) were trained using individual features (Leave P out, p=n-1), and the significant importance of each feature was assessed. Attribution to a specific body part was visualized using a semantic lineage tracing approach whereby the transformation path of each feature was back-traced to the originating body part. Each path on the map was then weighted if significant features passed along it. The resulting tree can then be interpreted; for example, Fourier-transformed paw-joint distances indicate a rhythmical motion pattern.

**Figure S. 5:**
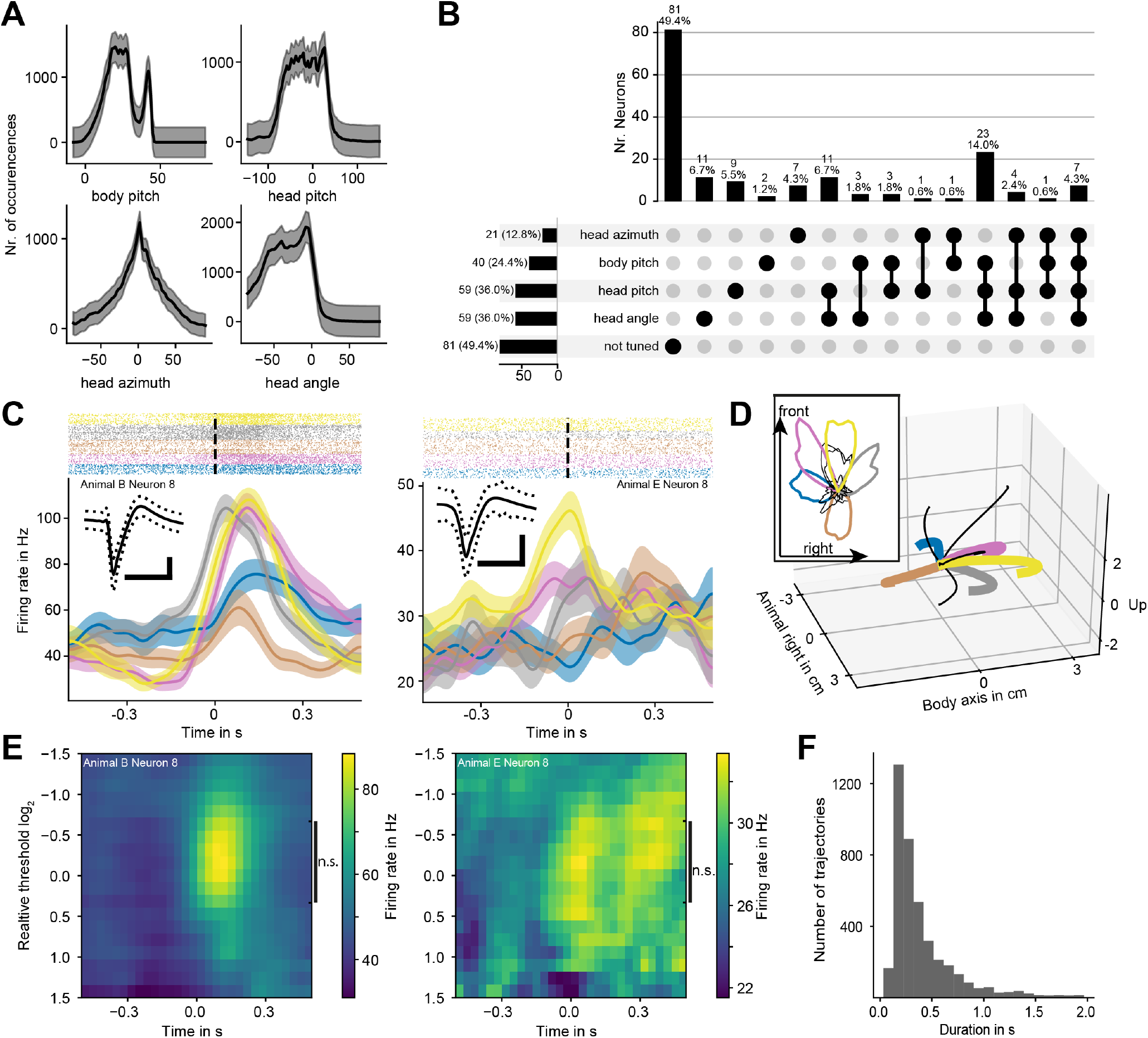
Posture and paw tuning multiplexing analysis. Related to Figure 4. **a** Histogram of the number of occurrences of data points in each bin for head and body pitch as well as head angle and azimuth. Shown is the mean along sessions, and shaded areas indicate std. The sharp second peak in the body pitch distribution corresponds to the rearing of the rats. **b** UpSet plot visualizing intersections and aggregates of sets of neurons with specific body posture tuning. Numbers indicate the number of neurons in each set. Bars indicate the number of neurons in each aggregated set, indicated via connection lines. On the left: aggregates across rows, top: aggregates across columns. Most tuned neurons coded for multiple postural variables. **c** Further example of neurons with a modulation of the neuronal response relative to the right-paw movements. Time point 0 represents the movement onset defined by a velocity threshold. Colors indicate the trajectory direction class and shaded areas indicate SEM. Horizontal bar indicates 1 ms and vertical 50 *µ*V. Inserts show the waveforms of the corresponding units ±STD. **d** Trajectories of the right (contralateral) paw. Mean trajectories for each class. Color represents the investigated four paw directions in the rat’s body reference frame. Inserts represent the distributions of the horizontal projections of movement directions. Colors indicate individual clusters defined by k-means. Some clusters were excluded from further analysis based on their vertical movement component (as opposed to the horizontal movement clusters) as well as consistency in trajectory length and direction. These clusters are represented by black lines. **e** Heat map of the neuronal response to paw motion using different thresholds. No significant difference in detected response was detected for all neurons when the threshold was varied between log_2_(−0.66) to log_2_(0.33) relative to the default threshold. One-way ANOVA of the sum of spikes within ± [333 ms] window around movement onset, with threshold as the main effect, per neuron Bonferroni-adjusted p*>* 0.3. **f** Distribution of paw-movement duration defined as time from threshold crossing to time point where the velocity decayed to 1/3 of its maximum.

**Figure S. 6:**
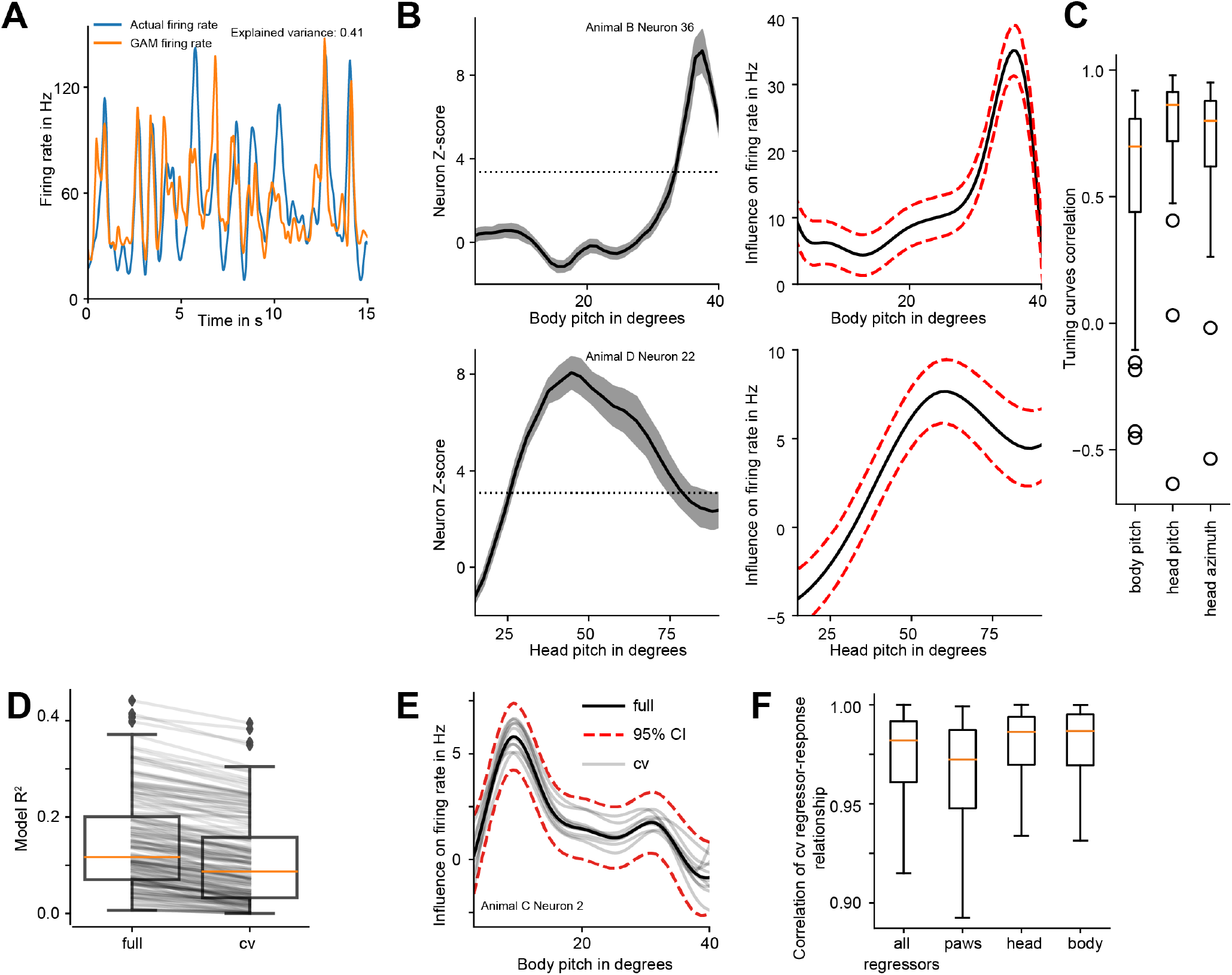
GAM analysis. Related to Figure 4. **a** Example of the actual firing rate and the GAM prediction for a neuron during a short representative period. **b** Comparison of the tuning predicted via GAM (right) with the tuning obtained using the bootstrap approach (left). Red dashed lines represent 95% confidence intervals (CI) of the model prediction. Black dashed lines indicate the significance threshold for the z-score, and shaded areas indicate SEM. **c** Comparison between tuning curves obtained via GAM and bootstrap approach indicates strong similarity. Correlations of the tuning between both methods for each neuron are shown as a boxplot. The box indicates the first and third quartiles. The median is represented by a horizontal line. Whiskers represent 1.5x IQR. **d** The model performance was similar, whether GAMs were fitted on full-data or cross-validated (cv) with part of the data used for training and part for testing. **e** Comparison of tuning predicted via GAM using the full data, or cv-models using parts of the data. **f** Tuning curves obtained via cv-GAMs are highly similar to the tunings curves obtained with the full-data models.

**Figure S. 7:**
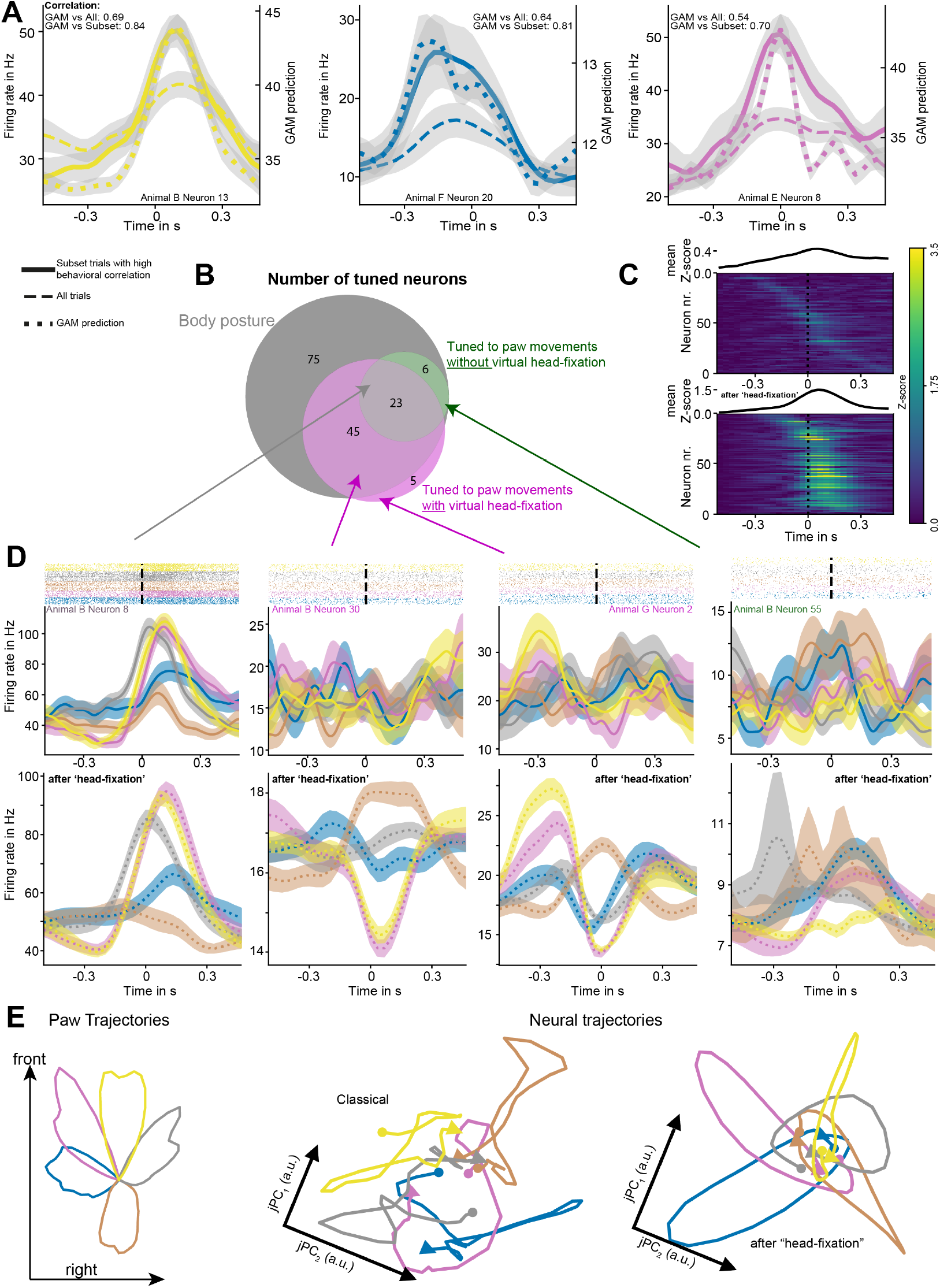
Supplement for “Virtual head-fixation”. Related to Figure 5. **a** Further individual examples for neuronal response to particular trajectory classes. Here, we contrast the average across trials from the entire session (dashed line) with the subset of paw movements with low postural variability (filled line). The GAM “head-fixed” prediction for the subset is shown as a dotted line. Inserts indicate the correlations between the individual GAM “head-fixed” predictions vs. subset or entire session. **b** Venn diagram displaying number of neurons detected as tuned for paw motion using both “classical” and “head-fixed” approaches as well as posture tuned as identified via GAM. Please note that the six neurons not detected using the “head-fixed’ approach were likely due to the strong contribution of potentially correlated body posture tuning. **c** Population response, revealed via “classic” (upper) or “virtual head-fixation” (lower) approaches. Neurons were sorted according to the time point of the maximal z-score within the response window. The “virtual head-fixation” approach was able to reveal much stronger responses in a larger fraction of neurons. Only neurons where GAM achieved explained variance *>* 0.1 are shown. **d** Further example neurons after “head-fixation”. Measured response to the trajectory class versus isolated predicted response following “virtual head-fixation”. Measured neural response is shown as filled line and “head-fixed” prediction as dotted line. **e** To visualize the population activity of neurons, we employed the jPCA technique(Churchland et al., 2012). After projection onto the first two components, the virtually headfixated population showed smooth neural trajectories (right), resembling the directionality of the paw trajectories (left). Without virtual head-fixation (middle), the trajectories also reflected movement directions but with a lower resemblance to that typically observed in head-fixed animals. Circles indicate trial start; arrows indicate trial end. Colors represent corresponding trajectory classes, and shaded areas represent SEM.

**Figure S. 8:**
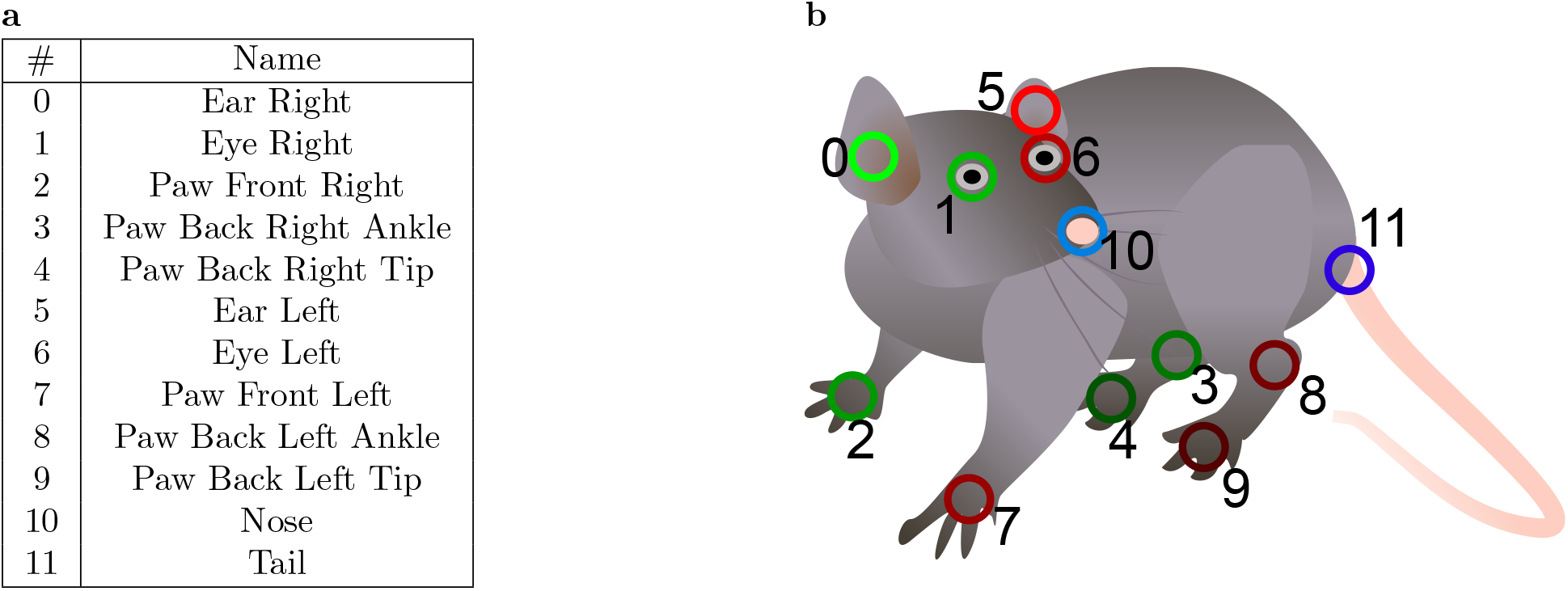
Keypoints defined on the rat body. Related to Fig1. **a** Names and indices of keypoints. **B** Keypoint locations on the rat body.

**Figure S. 9:**
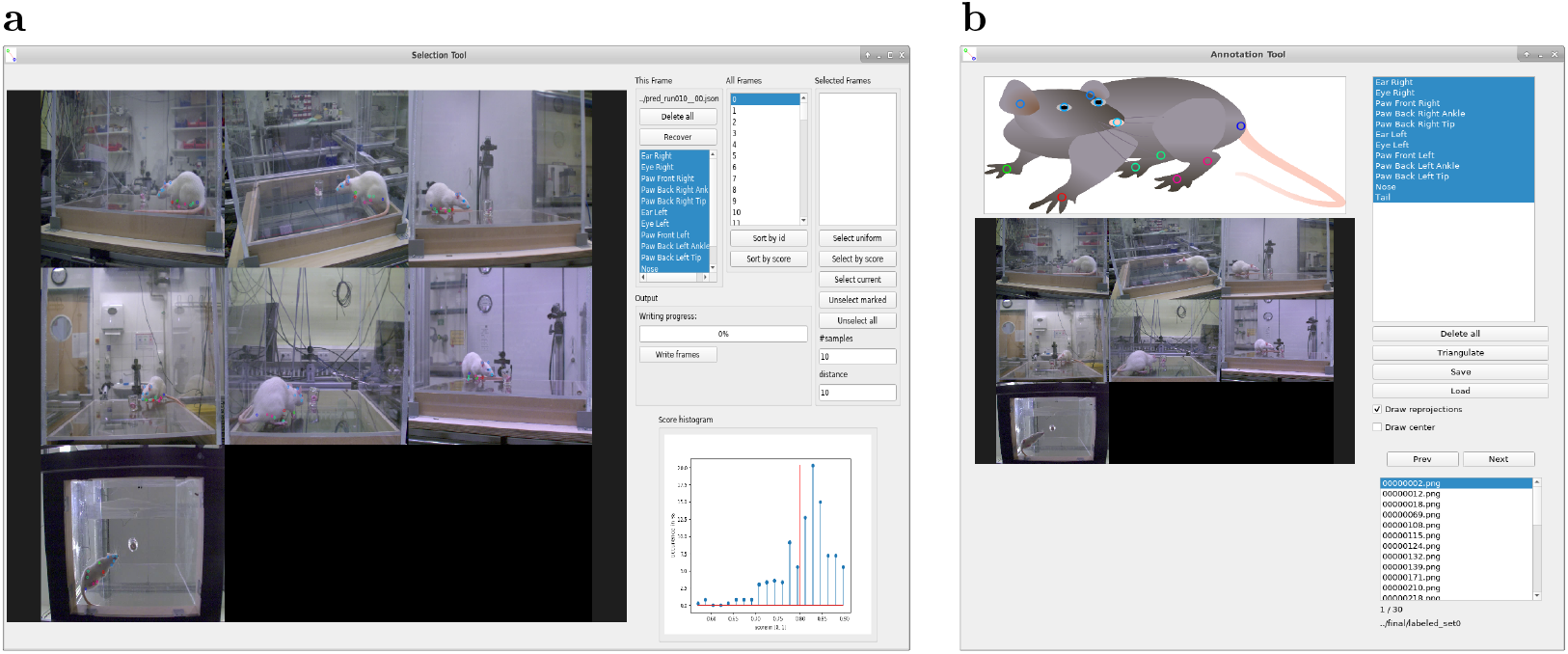
GUIs for viewing predictions and refining annotations. Related to Figure 1. **a** User interface for inspection and selection of frames, where refined annotation is necessary. The selection is guided by a confidence score of the network (lower right corner). **b** The selected frames are extracted from the videos and can be labeled by the annotation tool, which enables the user to annotate by dragging keypoints from the example image and dropping them onto the camera frames. This leverages the multi-view information by calculating a 3D point hypothesis based on the annotated 2D points.

### GAM experiment on toy data

To visualize the GAM approach, which can be used to separate a combined influence of multiple predictor variables, we designed a toy example. In our case, we have a set of predictor variables, of which we do not know which subset, if any, is contributing to the response variable, i.e., the neural firing rate. In our simplified toy example, let there be 4 predictor variables *x*_1_, *x*_2_, *x*_3_, *x*_4_. We model a response variable y via

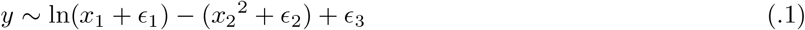

with *E* being Gaussian noise (Fig. S. 10**a**,**b**). We then fit our predictor variables via a step-wise forward-search procedure to the response variable y (Fig. S. 10**c**). The GAM approach was able to correctly reject the variables *x*_3_, *x*_4_ as well as correctly identify the individual contribution of the variables *x*_1_, *x*_2_ (Fig. S. 10**d**,**e**). The individual contribution is analogous to the “virtual head-fixation” approach: one variable keeps its values, whereas all other predictors are held constant.

**Figure S. 10:**
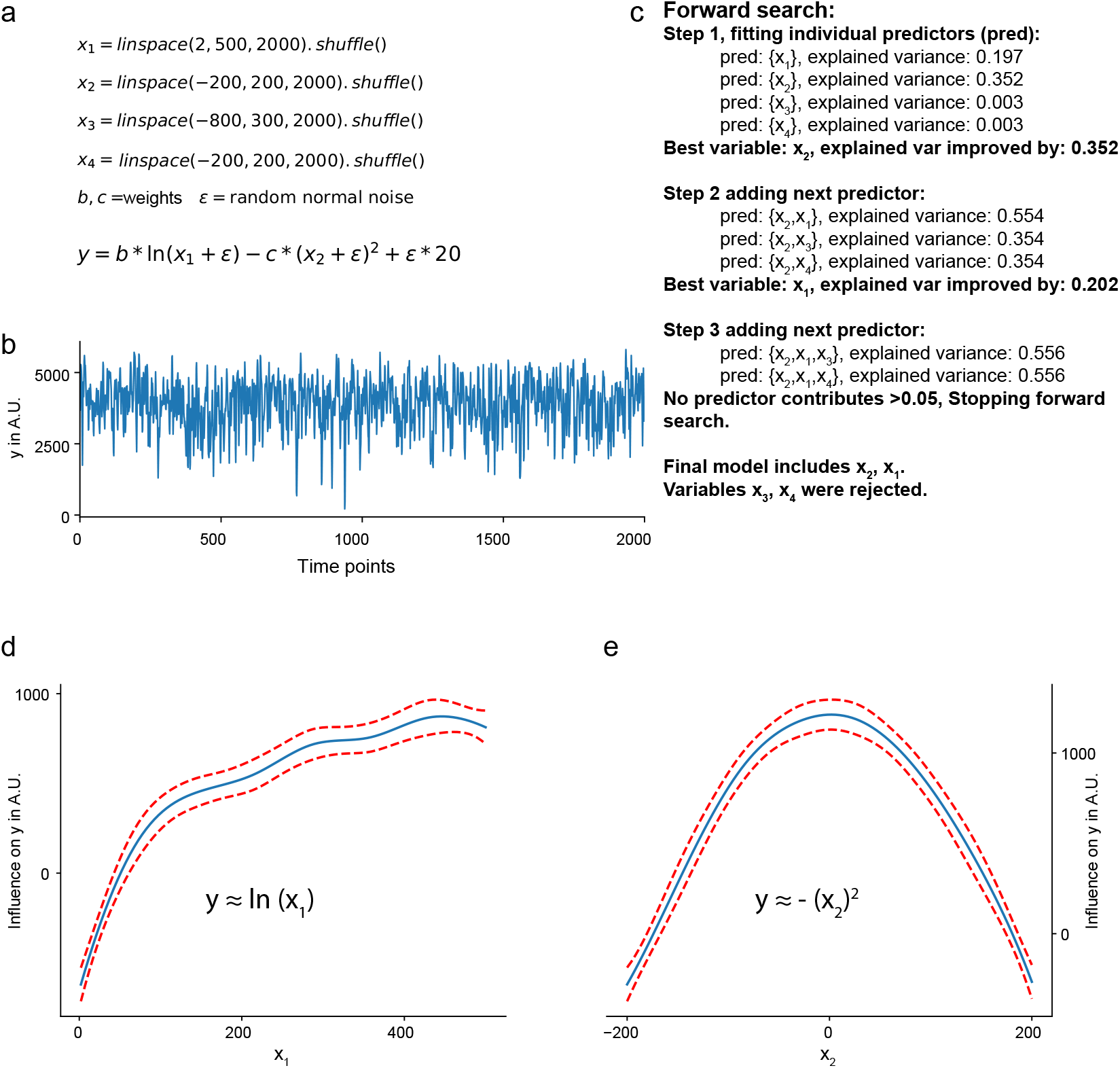
GAM toy example. Related to Figure 4d. **a** Generation of the predictor variables as well as the response variable. **b** Plot of the response variable y. **c** Step-wise, forward-search procedure. A model was fitted with each predictor, and only the best one is continuously added. **d** Predicted individual contribution of *x*_1_ on y showed a logarithmic relationship, corresponding to the expected relationship *y* ∼ ln(*x*_1_). **e** Predicted individual contribution of *x*_2_ on y showed a quadratic relationship, corresponding to the expected relationship *y* ∼ *x*_2_^2^. Red dashed lines represent 95% confidence intervals.

### FreiPose framework

We introduce a ready-to-use framework for general-purpose 3D keypoint estimation from multiple views, which is outlined in Fig. 1**c**. This framework is versatile in terms of the number of cameras and their relative positioning, as well as the number and semantic meaning of keypoints, which means it can be applied to a broad range of experimental setups.

The framework is easy to install because a *Docker* container is provided in our GitHub repository (https://github.com/lmb-freiburg/FreiPose-docker) along with video tutorials that provide a beginner-level introduction to how to use FreiPose for experimentation. We strongly recommend making use of these video tutorials.

We divided our framework into three modules, each of which can be used independently. The modules are RecordTool, FreiCalib, and FreiPose.

- **RecordTool** contains software for recording time-synchronized videos and directly handles the Basler cameras we used in our experiments. When using other cameras, this software must be modified accordingly.
- **FreiCalib** calculates the camera calibration based on time-synchronized video recordings, where a calibration object is used. Notes on how to create such a pattern and hints regarding what makes a good calibration sequence are provided in our respective GitHub repository.
- **FreiPose** estimates poses based on time-synchronized videos taken by previously calibrated cameras.

### Computational requirements

Running FreiPose involves training large neural networks so that particular hardware requirements should be met. This involves an Ubuntu 18.04 operating system and a nVIDIA GPU with at least 8GB of memory. Similar operating systems that also provide nVIDIA Docker might work as well but were not tested. Access to a solid-state drive (SSD) for storing training data is beneficial but optional.

### Video Recording with RecordTool

The module RecordTool builds upon the proprietary camera drivers of the Basler USB cameras that we used in our study and provides basic functions for camera usage, such as starting or stopping during recordings as well as modifying relevant camera parameters, including white balancing, frame rate, gain, and exposure time. Additionally, the same program operates the Arduino-based trigger system.

We designed RecordTool such that it saves the files into one output folder. In this folder, each recording is referred to as a *run* and consists of as many files as there are cameras. The files follow the naming convention `run%03d_cam%d.avi`, e.g. `run000_cam1.avi`. The software is available online alongside a detailed description.

### Camera Calibration with FreiCalib

First, video recordings using a suitable calibration object should be obtained. FreiCalib enables one to create a PDF file that can be used to manufacture a pattern calling

~~~
         python create_marker.py --tfam 36h11 --nx 10 --ny 4
                                 --size 0.05 --double
~~~

which will create two PDF files containing the front and back side of a pattern showing a 10 by 4 grid of tags (Olson, 2011) with size 5 cm each. Alongside the PDFs, a calibration pattern description file is saved that encapsulates marker information for subsequent processing steps.

For a given recorded calibration sequence, the important parameters describing the imaging process can be calculated by running

~~~
    python calib_M.py {MARKER_PATH} {VID_PATH}
~~~

where `{MARKER_PATH}` is the path to the marker description file, and `{VID_PATH}` points to the videos showing the calibration sequence. The calibration result will be saved to the folder of the videos as M.json. The calibration file contains readable characters and can be opened with any text editor. This file contains a dictionary of lists, where each dictionary item presents either extrinsic, intrinsic, or distortion parameters, and the list iterates over cameras.

Additionally, we provide the option to check a previously calculated calibration with respect to a newly recorded video sequence.

~~~
     python check_M.py {MARKER_PATH} {VID_PATH} {CALIB_FILE}
~~~

This option enables easy verification of the given calibration’s validity, e. g.none of the cameras was (accidentally) moved or altered between calibration and experimentation time.

**Table S. 1:** Camera subsets for reduced number of views experiment. Given a number of cameras, three different subsets of cameras are selected. Reducing the number of cameras leaves both fewer frames for training and increases the difficulty to precisely localize keypoints (Fig. 1**e**). Please note that extensive evaluation of all possible configurations is computationally very expensive, and we therefore resort to using manually selected subsets that reflect reasonable camera placements.

| Number of cameras | 1 | 2 | 3 | 4 | 5 | 6 |
| --- | --- | --- | --- | --- | --- | --- |
| Camera sets | {1} | {1, 5} | {1, 5, 7} | {1, 3, 4, 5} | {1, 3, 4, 5, 7} | {1, 2, 3, 4, 5, 7} |
|  | {5} | {1, 7} | {1, 3, 5} | {1, 3, 4, 7} | {2, 6, 4, 5, 7} | {1, 2, 3, 4, 5, 6} |
|  | {7} | {1, 3} | {4, 5, 7} | {1, 4, 5, 7} | {1, 3, 4, 5, 6} | {1, 2, 4, 5, 6, 7} |

### Experimentation using FreiPose

After obtaining a valid calibration, the actual experiment can be conducted. To begin, an initial configuration of the framework is necessary, which mainly involves defining the keypoints required for the task and additional task-specific settings. A detailed description of these parameters is located in the GitHub repository and is encapsulated in a configuration file denoted by `{CFG_FILE}`.

To predict poses on a given set of videos, the calls

~~~
   python predict_bb.py {CFG_FILE} {VID_PATH}
   python predict_pose.py {CFG_FILE} {VID_PATH}
~~~

are sufficient. The commands will save predictions into a new file denoted by `{PRED_FILE}` in the folder of the videos, and its content can be visualized with

~~~
   python show_pred.py {CFG_FILE} {PRED_FILE}
~~~

To inspect the predictions in detail and select frames for manual annotation, the Selection Tool (Fig. S. 9**a**) is used. This tool is started by calling

~~~
   python select.py {CFG_FILE} {PRED_FILE}
~~~

The user is guided through the selection process by prediction confidences and automatic selection methods presented by the GUI. Frames selected for labeling are extracted from the videos and saved in a separate folder `{FRAME_FOLDER}` as individual frames. The labeling tool (Fig. S. 9**b**) can be used in parallel by multiple persons and is started by

~~~
   python label.py {CFG_FILE} {FRAME_FOLDER}
~~~

The labeling tool can load the networks’ pose predictions, which accelerates the labeling procedure. As the networks’ predictions improve, the user can resort to correcting erroneous labels instead of labeling the complete set of keypoints from scratch. The labeling tool leverages multi-view geometry (i.e., it triangulates the annotations in the individual camera views to a 3D hypothesis and visualizes their locations). This approach allows the annotator to stop annotating as soon as the hypothesis is consistent across views, which yields a tremendous reduction of labeling effort because the annotation in a few camera views is typically sufficient.

The folder containing the labeled data is registered to the framework by creating an entry in its configuration file. Afterwards, the network can be fine-tuned by calling

~~~
   python train_bb.py {CFG_FILE}
   python train_pose.py {CFG_FILE}
~~~

### Details of the Motion Capture Experiment

The experimental motion capture results of 2 were obtained by splitting the base dataset of 1199 training and 614 evaluation frames providing 7 cameras into different subsets. For a given number of cameras, the experiment is repeated in 3 trials choosing different sets of cameras as listed in Table S. 1. For example, if only 2 cameras are used, we select the following camera pairs: {{1, 5}, {1, 7}, {1, 3}}. Each number uniquely identifies a camera (Fig. 1**e**), and the pairs chosen correspond to the cases “long side + short side”, “long side + bottom”, and “long side + long side”. Each of the resulting datasets still covers 1199 time instances but only 1199 · 2 = 2398 individual frames compared to 1199·7 = 8393 in the all-camera setting. The same procedure is applied to the evaluation set. Table S. 1 lists the selected subsets of cameras used for experiments in Fig. 1**e**. Please note that testing all possible permutations is computationally very expensive; therefore, we resort to testing manually chosen subsets representing meaningful cases, i. e. chose cameras how one would if only a limited number of cameras is available.

To simulate the sparsity of labeled samples (Fig. 1**e**), we use all cameras but randomly select a subset of 10%, 20%, 30%, 60%, 80% or all time instances. For example, in the 20% case, there are 1199·0.2 = 239 time instances of 7 cameras in the training set, which results in an effective number of 239·7 = 1673 camera frames used. This dataset is used for training both methods, FreiPose and DLC, and the evaluation is performed with respect to the complete evaluation set. Each level of sparsity is sampled 3 times for a more robust reconstruction.

### Inference speed

For most applications of post hoc motion tracing, inference speed is only of secondary priority. Thus, FreiPose was designed with accuracy, not speed, as its priority. However, we still analyzed the inference speed and performed some optimizations towards the real-time inference goal.

To this end, we first investigated the speed at which each part of the algorithm runs. We identified the Pose Estimation node as a major bottleneck and targeted it for modifications. First, we pruned the 2D encoder network so that it uses fewer channels for feature representation (“2D slim”)—a strategy that resulted in a large loss in pose estimation accuracy while only yielding a modest increase in processing speed. Second, we decreased the spatial resolution of the 3D grid that the 3D CNN is operating in. The base version used 64 voxels per dimension, whereas the tested variants utilized 32 (“3D V32”) and 16 (“3D V16”), respectively. Cutting the voxel resolution in half resulted in a large acceleration (186% faster), whereas only a marginal increase in mean position error was observed (11.0% larger). Reducing the resolution even further to 16 led to an even larger improvement in terms of FPS (261.3% faster) but induced a much worse pose error (45.6% larger).

| Variant | base | 2D slim | 3D V32 | 3D V16 |
| --- | --- | --- | --- | --- |
| FPS | 3.63 | 4.25 | 9.89 | 12.41 |
| Mean Per Joint Position Error | 7.09 | 13.46 | 7.87 | 10.32 |

### Optimization of the pose node

Largest potential lies in optimization of the 3D part of the architecture, whereas modifying the 2D part only yields relatively small increases in speed at large cost in terms of pose accuracy.

Even more room for improvement was identified in the hardware used. The compute unit, currently in use, had a Titan X, but much faster GPUs are readily available. Table lists different runtimes achievable on different GPU models and showed a substantial increase changing nothing but one piece of hardware, which approximately doubles the number of frames processable by the node.

| GPU | Titan X | 1080 | 1080Ti | 2080Ti |
| --- | --- | --- | --- | --- |
| FPS | 9.89 | 11.01 | 15.00 | 19.98 |

### Runtime of the pose node dependent of GPU model used for computation

A large dependency towards the hardware used was revealed. From slowest to fastest GPU lies a factor of 2, which partially turns the question of runtime into simply using the correct compute platform.

Using the optimized network architecture and the more recent 2080Ti GPU, the system shows a total runtime of 0.126 s, equivalent to a processing speed of 7.94 Hz for the complete system. With further improvements in hardware available and/or usage of multi-GPU arrays, real-time applications might become feasible soon. However, this was not the main focus of our work because our analysis did not require real-time evaluation. Thus, throughout the paper, we used the most accurate ”base” version of FreiPose.

